# Genetic and functional diversification of chemosensory pathway receptors in mosquito-borne filarial nematodes

**DOI:** 10.1101/683060

**Authors:** Nicolas J Wheeler, Zachary W Heimark, Paul M Airs, Alexis Mann, Lyric C Bartholomay, Mostafa Zamanian

## Abstract

Lymphatic filariasis (LF) afflicts over 60 million people worldwide and leads to severe pathological outcomes in chronic cases. The nematode parasites (Nematoda: Filarioidea) that cause LF require both arthropod (mosquito) intermediate hosts and mammalian definitive hosts for their propagation. The invasion and migration of filarial worms through host tissues are complex and critical to survival, yet little is known about the receptors and signaling pathways that mediate directed migration in these medically important species. In order to better understand the role of chemosensory signaling in filarial worm taxis, we employ comparative genomics, transcriptomics, reverse genetics, and chemical approaches to identify putative chemosensory receptor proteins and perturb chemotaxis phenotypes in filarial worms. We find that chemoreceptor family size is correlated with the presence of environmental (extra-host) stages in nematode life cycles, and that filarial worms contain a compact and highly-diverged chemoreceptor complement and lineage-specific ion channels that are predicted to operate downstream of chemoreceptor activation. In *Brugia malayi*, an etiological agent of LF, chemoreceptor expression patterns correspond to distinct parasite migration events across the life cycle. To interrogate the role of chemosensation in the migration of larval worms, arthropod infectious stage (microfilariae) and mammalian infectious stage (L3) *Brugia* parasites were incubated in nicotinamide, an agonist of the nematode transient receptor potential (TRP) channel OSM-9. Exposure of microfilariae to nicotinamide alters intra-mosquito migration while exposure of L3s reduces chemotaxis towards host-associated cues *in vitro*. Nicotinamide also potently modulates thermosensory responses in L3s, suggesting a polymodal sensory role for *Brugia osm-9*. Reverse genetic studies implicate both *Brugia osm-9* and the cyclic nucleotide-gated (CNG) channel subunit *tax-4* in larval chemotaxis towards host serum, and these ion channel subunits rescue sensory defects in *C. elegans osm-9* and *tax-4* knock-out strains. Together, these data reveal genetic and functional diversification of chemosensory signaling proteins in filarial worms, and encourage a more thorough investigation of clade and parasite-specific facets of nematode sensory receptor biology.

## Introduction

Lymphatic filariasis (LF) is a parasitic disease caused by mosquito-borne filarial worms (Nematoda: Filarioidea) belonging to the genera *Wuchereria* and *Brugia*. LF is estimated to affect over 60 million people worldwide, particularly in impoverished tropical regions (1). Infections are associated with chronic disability and physical disfigurement, most commonly resulting from advanced manifestations of lymphedema, hydrocele, and elephantiasis. These gross manifestations yield additional stigmatisation and mental health burdens on those suffering, which in turn can prevent individuals from seeking treatment (2–4). Currently, chemotherapeutic control of LF is mainly achieved through mass drug administration (MDA) of diethylcarbamazine citrate (DEC), ivermectin (IVM), albendazole, or combinations of these anthelmintic drugs (5,6). However, the suboptimal efficacy of available drugs against adult parasites, contraindication of DEC and IVM in patients with multiple filarial diseases, and the threat of drug resistance underlie efforts to develop new treatment options. A better understanding of the molecular basis of parasite behaviors required for successful transmission and parasitism has the potential to aid LF control efforts.

The filarial worms that cause LF have complex life cycles that require migration through hematophagous arthropod intermediate hosts and vertebrate definitive hosts (7). Microfilariae (mf) released from viviparous females in the human lymphatics must reach the peripheral blood where they can be taken up by the proboscis of feeding mosquito intermediate hosts. In susceptible mosquitoes, larvae burrow out of the mosquito midgut, pass through the hemocoel and invade cells of thoracic flight muscles. Larvae grow and develop over the course of approximately two weeks to the human-infective third larval stage (L3), which migrate to the mosquito head region in preparation for transmission to the mammalian host (8,9). L3s are deposited onto the skin of vertebrate hosts from the proboscis of feeding mosquitoes and must quickly travel through the bite wound and connective tissues to reach the lymphatic system where they reach sexual maturity (10,11). While the life cycle of LF parasites are well described, the molecular basis for stage-specific migratory behaviors is unknown.

There is growing evidence that chemosensory and other sensory modalities play an important role in nematode parasite transmission and intra-host migration (12–21). However, most studies have focused on single-host nematode parasites with direct life cycles, which are phylogenetically distant from the vector-borne filarial parasites of clade III. Recent studies using human-infective *Brugia malayi* and feline-infective *Brugia pahangi*, a model species for human LF, reveal the presence of canonical nematode sensory organs (amphids) and robust chemotaxis responses to host-associated cues *in vitro (22–25*). Filarial worms also exhibit genus-specific patterns of intra-host migration (26). These observations strongly suggest an important role for chemosensation and chemotaxis in LF parasitism and provide motivation to dissect the signaling pathways and mediators of sensory behaviors in these medically important parasites.

Chemosensory signaling pathways in the model nematode *Caenorhabditis elegans* are well-characterized (27). G protein-coupled receptors (GPCRs) function as chemoreceptors at the amphid cilia, and activation leads to signaling through cyclic nucleotide-gated channels (CNGs) or transient receptor potential channels (TRPs), depending on cell type (28–31). Each amphid neuron expresses a diverse array of GPCRs, in contrast to the one-receptor-per-cell model in vertebrates (32–34). These pathways have likely evolved to reflect the diversity of nematode life-history traits and environmental cues encountered by different nematode species (13,18–20). Despite superficial conservation of nematode chemosensory pathways, we hypothesized that there are important differences in chemosensory gene repertoire, patterns of expression, and function among free-living, single-host, and vector-borne parasitic nematodes belonging to diverse clades (35,36).

Here, we investigate nematode chemosensory receptor biology in LF parasites and connect *in vitro* and *in vivo* chemotaxis behaviors to chemosensory signaling pathways. We carry out genomic and transcriptomic analyses of putative chemosensory GPCRs (chemoreceptors), CNGs, and TRPs in a pan-phylum context. Using a combination of chemical and reverse genetic approaches, we present the first evidence of *Brugia* chemotaxis behaviors modulated by specific sensory-associated receptors. Lastly, we explore how these data reveal unique aspects of chemosensory biology in these medically important parasites.

## Results

### Filarial worms contain a compact and unique repertoire of chemoreceptors

To elucidate the putative chemosensory pathway of mosquito-borne filarial worms and to identify and annotate chemoreceptors, we first performed a pan-phylum analysis of 39 nematode genomes (37,38), representing all published filarial genomes and high-quality assemblies across four primary nematode clades (36) (S1 Table, S1 Figure). 10,440 chemoreceptor genes were identified and confidently classified within superfamilies (Str, Sra, Srg) or “solo” families (srz, sro, srsx, srbc, srxa, sra) (S1 File) (39). While the majority of receptors were also annotated at the family level, some clade IIIa/b and clade IV chemoreceptors did not clearly group with the families that were originally described in *C. elegans* (Fig 1B-C, S1 Data). However, each of the 30-100 chemoreceptors found in filarial worm species (clade IIIc) were readily classified into the established 23 nematode chemoreceptor families (39,40). Within these families we found no one-to-one orthologs between filarial parasites and species belonging to other clades, demonstrating the divergence of the filarial chemoreceptor subset. Instead, there have been clear paralogous gene radiations that have resulted in enrichment of the *srx, srab, srbc*, and *srsx* families (Fig 1B-C). Filarial parasites also contain relatively numerous *srw* receptors, but these likely include neuropeptide receptors in addition to some chemoreceptors of environmental peptides (40,41).

**Figure 1.**
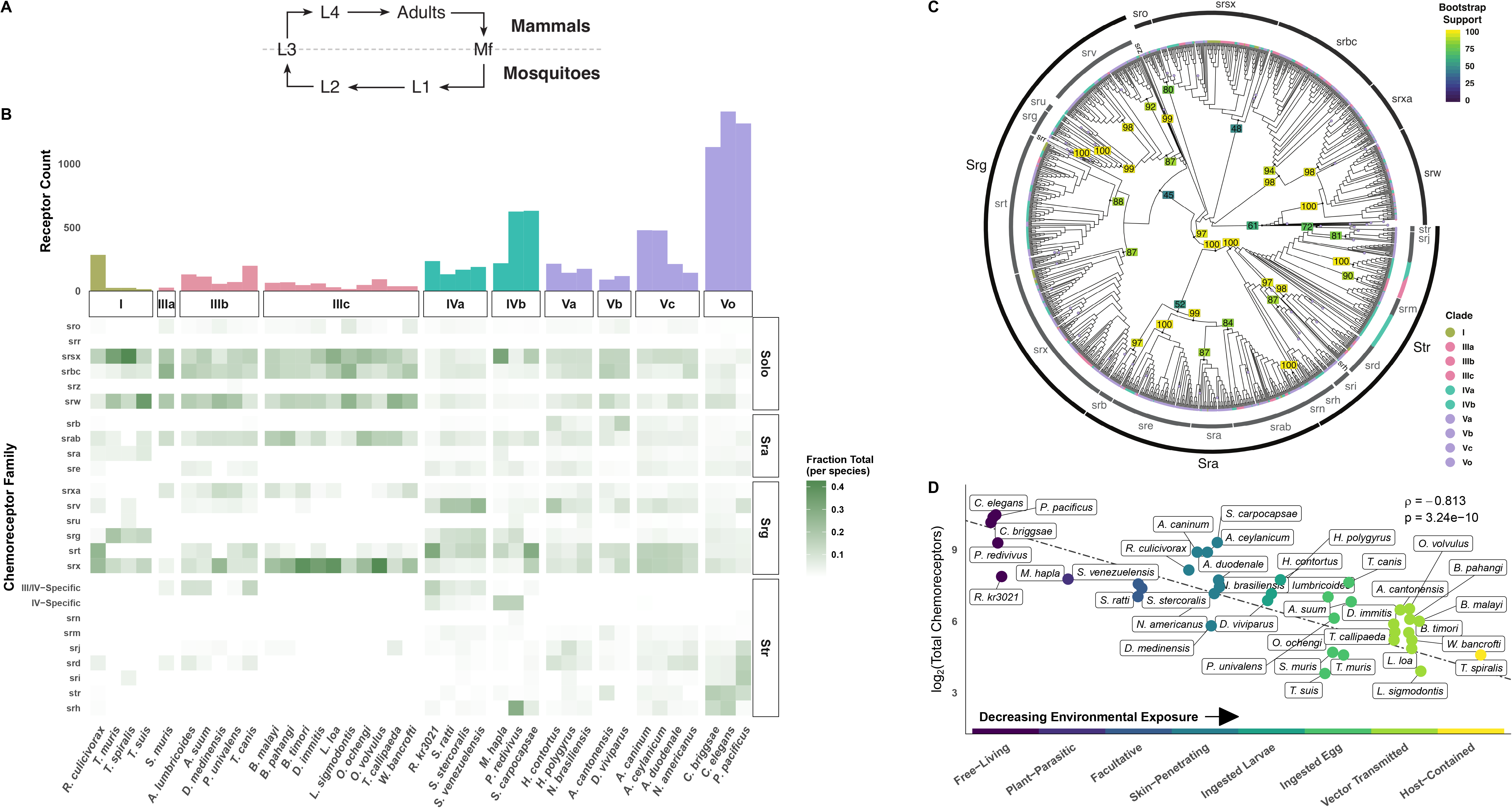
The genomes of filarial worms contain a reduced complement of divergent chemoreceptors. Chemoreceptors were mined from 39 nematode genomes, and the phylogeny of chemoreceptors from a down-sampled species set was constructed with maximum-likelihood (ML) inference. **(A)** General life cycle patterns of mosquito-infective filarial worms. **(B)** Family and superfamily categorizations were used to annotate the final phylogeny. Clade IIIc chemoreceptors are diverged from *C. elegans* and other nematode chemoreceptors, without any one-to-one homologs. Filarial chemoreceptors are notably diverged in *srsx, srab*, and *srx*. Nodal values represent bootstrap support of 1000 separate bootstrap replicates. Branches consisting of only *C. elegans* receptors were collapsed to aid visualization. **(C)** Filarial worm genomes contain far fewer chemoreceptors than other parasitic and free-living nematodes, and they are enriched for *srsx, srab*, and *srx* receptors. Each box in the heatmap is normalized to the total number of chemoreceptors per species. Values from the bar plot in B are log2 transformed. **(D)** A decrease in chemoreceptor count is correlated with an increase in extra-host (e.g., terrestrial) stages within nematode life cycles. Completely free-living nematodes such as *C. elegans, C. briggsae, P. pacificus*, and *P. redivivus* have many more chemoreceptors than parasitic nematodes that are vector-transmitted or host-contained such as the filarial worms and *T. spiralis*. Note that the x-axis is categorical and slight jitter has been added to the points to decrease point/label overlap and aid interpretation, ρ was calculated with Spearman’s rank correlation with the null hypothesis that ρ = 0. See https://zamanianlab.shinyapps.io/ChemoR/ for an interactive version of (D).

Filarial worm genomes contain a reduced subset of chemoreceptors when compared to other parasitic and free-living nematodes, including *C. elegans* and *C. briggsae*, both of which contain over 1200 chemoreceptors (clade V) (Fig 1B) (34,39,40). While it is known that parasitic nematodes contain fewer chemoreceptor genes than *C. elegans* (42,43), and indeed often fewer genes in total(44), our pan-phylum analysis revealed a significant correlation between chemoreceptor gene count and the presence and nature of free-living or environmental stages of each nematode species life cycle (Figure 1D, S2 File). Parasites that are more host-contained and lack motile environmental stages exhibit more compact chemoreceptor repertoires than those that are exclusively free-living or contain free-living stages (Spearman’s rank-order correlation, ρ = −0.813, p = 3.24 x 10^−10^), and this is not a function of genome contiguity or completeness (S2 Figure). This correlation has been observed when comparing smaller numbers of nematodes, and our comprehensive approach confirms this pattern across the phylum (39,43,45). An interactive version of Fig 1 and S2 Figure is available at https://zamanianlab.shinyapps.io/ChemoR/. where the annotation and amino acid sequence data for the chemoreceptors in all 39 nematode genomes is available for download.

Chromosomal synteny between *Brugia malayi* and *C. elegans* further illustrates the divergence of chemoreceptors in the Filarioidea (Fig 2). The majority of *C. elegans* chemoreceptors are found on chromosome V (67%) and likely underwent several birth-death cycles that reflect the adaptive needs of encountering new locales (39). Putative *B. malayi* chemoreceptors are primarily found on two chromosomes, II (31%) and IV (35%), and are clustered by family, suggesting lineage-specific gene duplications.

**Figure 2.**
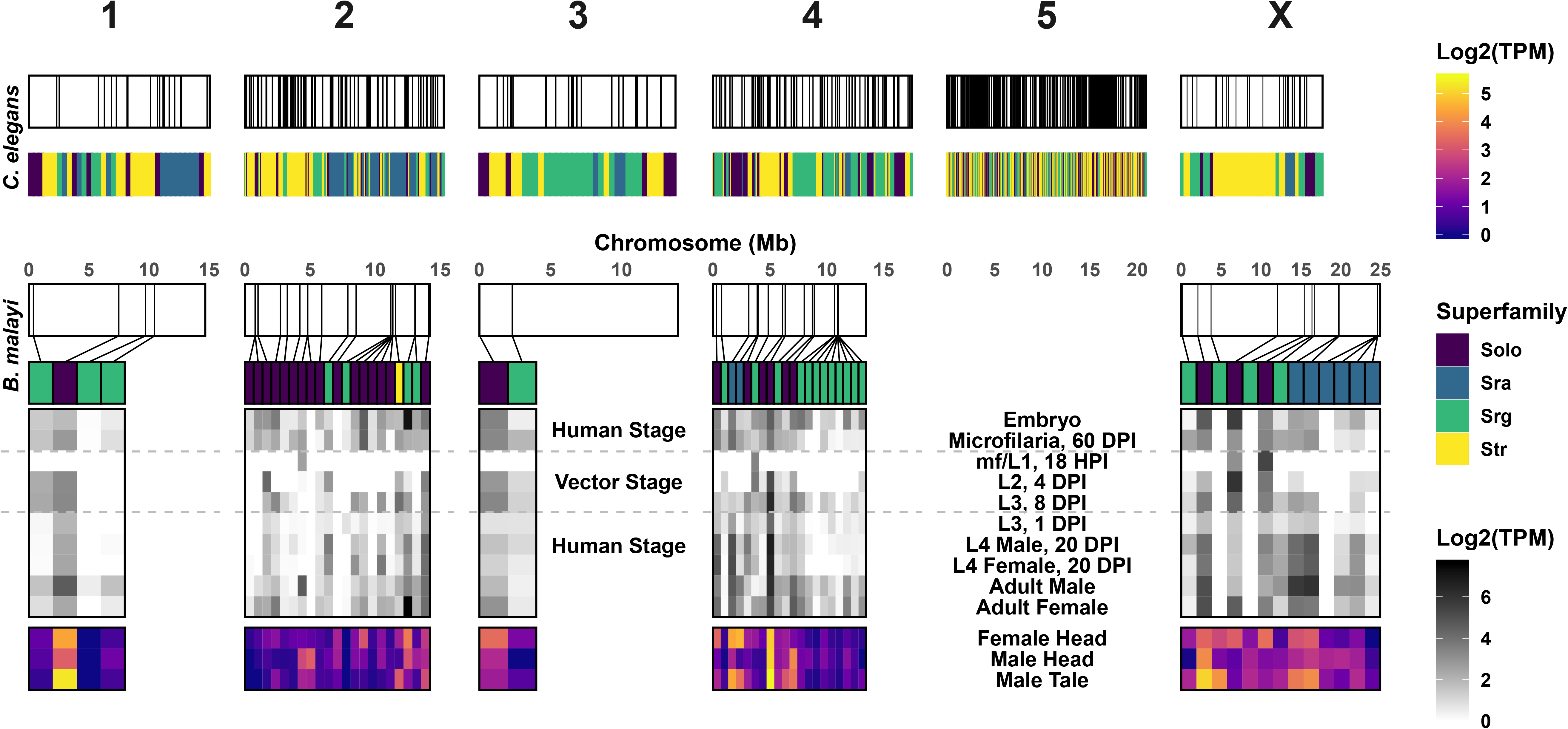
Chemoreceptors are clustered in the *Brugia malayi* genome and are enriched in specific life stages and adult tissues. The chromosomal location of annotated *B. malayi* and *C. elegans* chemoreceptor genes are shown with chromosomes in white and chemoreceptor loci as black lines. *C. elegans* chemoreceptors are found throughout the genome but are heavily clustered on chromosome V, and these clusters can be enriched for specific families and superfamilies (39). Likewise, *B. malayi* chemoreceptors are clustered on chromosomes II and IV. RNA expression data reveal distinct patterns of chemoreceptor expression across the life cycle and in discrete adult male and female tissues.

### Brugia malayi *chemoreceptors are associated with sensory tissues and display stage-specific expression patterns*

These comparative data indicate that arthropod-borne filarial worms rely on a small complement of clade and species-specific chemoreceptors to interact with and navigate their host environments. Nematode chemosensation is primarily mediated by anterior amphid sensory structures, and many nematodes also possess caudal chemosensory phasmids associated with male sex organs and likely aid in copulation. In *C. elegans* hermaphrodites and *Strongyloides stercoralis*, a soil-transmitted helminth, chemoreceptors and sensory pathway effectors have been primarily localized to these anterior and posterior structures but can also be found in other non-neuronal cells (12,34). We examined the expression of chemoreceptors in structures implicated in adult filarial worm chemosensation (22,46,47) using RNA-seq of anterior and posterior tissues. *B. malayi* female head, male head, and male tail tissue regions were excised for RNA-seq detection of anterior, posterior, and sex-specific chemoreceptor transcripts. Most chemoreceptor transcripts are preferentially detected in one of these disparate anatomical regions, while a small number show a broader distribution of expression across these regions (Fig 2, S3 Figure).

We further hypothesized that the unique cues encountered by filarial parasites across developmental time points would be reflected by stage-specific chemoreceptor expression patterns. In *C. elegans*, chemosensory processes coordinate movement towards food or mates and away from pathogens, predators, or noxious substances (48–51). In contrast to the open and less predictable environments navigated by free-living nematodes, filarial worms encounter distinct environmental niches that have strictly patterned transitions. We used staged transcriptomes to analyze the expression of chemoreceptors across the life cycle of *B. malayi* (52) and identified receptors that correspond to migratory landmarks throughout the parasite life cycle (Fig 2). Expression data show that microfilariae circulating in the bloodstream at 60 days post-infection (DPI) express a larger number of chemoreceptors compared to non-migratory L1 and L2 larvae that are contained within mosquito muscle cells; for instance, only 4 chemoreceptors during the mf/L1 stage in the mosquito have detectable expression (Fig 2). There is an increase in chemoreceptor representation and expression during the migratory and mammalian-infective L3 stage, as well as in later mammalian stages that undergo migration and potentially engage in mate-seeking behaviors. Together, these analyses show that *B. malayi* expresses distinct sets of chemoreceptors in a sex, tissue, and stage-specific manner.

### Filarial worms have a divergent subset of downstream chemosensory pathway receptors

In *C. elegans*, ligand binding to chemoreceptors activates heterotrimeric G proteins that ultimately produce neuronal depolarization via the opening of cyclic nucleotide-gated channels (CNGs) or transient receptor potential channels (TRPs), depending upon cell type (27,34). The CNGs TAX-4 and TAX-2 mediate signaling in amphid neurons ASE, AWC, AWB, ASI, ASG, ASJ, ASK, while the TRPV (vanilloid-type) channels OSM-9 and OCR-2 are necessary for signaling in AWA, ASH, ADF, and ADL (27). To assess the conservation of these downstream signaling pathways in filarial parasites, we mined TRP and CNG channels across nematode genomes to examine interspecies variation in ion channel complements.

We found that filarial worms contain one-to-one homologs of *osm-9*, but do not have homologs of *ocr-3, ocr-4, trpa-1, pkd-2, trp-1*, or *gtl-1* (Fig 3A/C/E, S2 Data). Filarial parasites contain two *ocr-1/2*-like genes (*Bm5691* and *Bm14098*), but these are more closely related to each other than they are to *C. elegans ocr-1* or *ocr-2* (Fig 3E). In *C. elegans*, OSM-9 and OCR-2 are mutually dependent upon each other for intracellular trafficking to sensory cilia (31). Cell-specific TRP channel expression patterns and TRP subunit interactions are unknown in filarial parasitic species, and it is not clear which filarial parasite subunit might provide a homologous Cel-OCR-2 ciliary targeting function, or indeed if such a trafficking function is necessary. Interestingly, we found that *Bma-ocr-1/2a* (*Bm5691*) is expressed in the female head (transcripts per million (TPM > 2.5) but is found in very low abundances in the male head and tails (transcripts per million (TPM) < 1), while the opposite is true for *Bma-ocr-1/2b* (*Bm14098*). On the other hand, *Bma-osm-9* is found at a relatively high abundance in both male (TPM > 15) and females (TPM > 18) heads. This tissue distribution of transcripts could indicate the potential for sex-specific subunit interactions among these TRPV channels. We found homologs of *ocr-3, ocr-4, pkd-2*, and *trp-1* in other clade III species (e.g. soil-transmitted ascarids), and the most parsimonious explanation of their absence in filarial worms is that these genes were lost sometime after the divergence of Spirurida and Ascarida (35). Conversely, *trpa-1*, which functions in *C. elegans* mechanosensation in QLQ (53), and *gtl-1*, which functions in ion homeostasis in the *C. elegans* intestine (54), appear to be specific to clade V.

**Figure 3.**
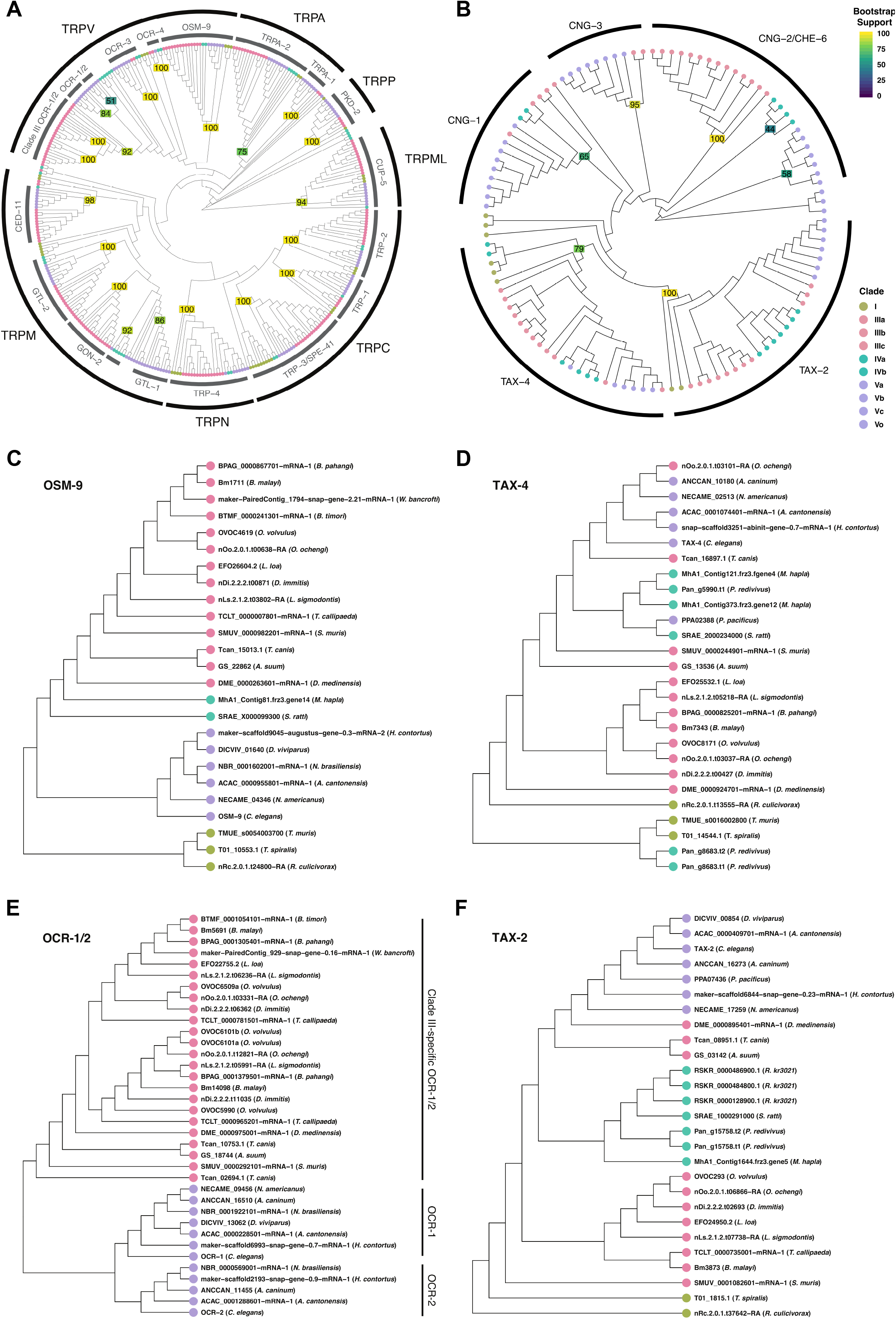
Filarial worms possess unique complements of broadly conserved nematode TRP and CNG channels. The phylogenies of **(A)** TRP and **(B)** CNG channels were constructed with Bayesian inference. Nodal values represent the posterior probability. **(C)** *osm-9* and **(E)** *ocr-1/2* subtrees were drawn from A, and **(D)** *tax-4* and **(F)** *tax-2* subtrees were drawn from B. Filarial worms have one-to-one orthologs of *C. elegans osm-9, tax-4*, and *tax-2*. In contrast, the two *ocr-1/2*-like genes from filarial worms are more closely related to each other than with the homologous *Cel-ocr-1* and *Cel-ocr-2*, and belong to a diverged clade IIIc grouping of OCR-1/2-like channel subunits.

Similarly, filarial worms have one-to-one homologs of *tax-4* (α-type) and *tax-2* (β-type) CNG channel subunits, but lack *cng-1* and *cng-3* (Fig 3B/D/F, S3 Data). Filarial worm genomes possess a third CNG (Bm7148) that is related to both *cng-2* and *che-6*, but phylogenetic analysis suggests the divergence of *cng-2* and *che-6* to have occurred later than the most recent common ancestor of the Filarioidea and *C. elegans*, making it difficult to ascribe putative function to the *cng-2/che-6* homolog in Filarioidea. In *C. elegans*, TAX-2 and TAX-4 are broadly expressed in amphid sensory neurons and mediate both thermosensory and chemosensory function, while other channels like CNG-2 and CHE-6 have more restricted expression patterns and modulate these pathways (55). It is unclear whether *Bma*-TAX-4 and *Bma*-TAX-2 coordinate multiple sensory modalities as in *C. elegans* and if *Bm7148* interacts with these proteins and pathways. *Bm7148* is highly expressed in all three tissues that we analyzed (TPM > 20) while *Bma-tax-4* and *Bma-tax-2* have TPM values of less than 2 in all cases.

### *Treatment with a nematode TRPV agonist inhibits chemoattraction but not chemoaversion of infective-stage* Brugia *larvae*

Our bioinformatic analyses show that filarial worms have evolved divergent sets of chemoreceptors, but maintain much of the core structure of the chemosensory pathway as modeled in *C. elegans*. To test conservation of chemosensory function in TRPV channels of filarial worms, we treated infective-stage *Brugia* L3s with nicotinamide (NAM), an agonist of the *C. elegans* OSM-9/OCR-4 heteromeric channel (56), and measured chemotactic responses to host-associated cues. These experiments were performed with *Brugia pahangi*, a model *Brugia* species (22,23,25). *B. pahangi* L3s freshly extracted from infected *Aedes aegypti* are strongly attracted to both fetal bovine serum (FBS) and sodium chloride but are weakly repelled by 3-methyl-1-butanol (a component of human sweat attractive to *Strongyloides stercoralis* and *Anopheles gambiae*) (8,13,49) (Fig 4B). Treatment of freshly extracted L3s with 250 μM NAM significantly reduced chemoattraction to serum (45.2% reduction) and sodium chloride (61.7% reduction), but had no significant effect on aversion to 3-methyl-1-butanol (Fig 4B). NAM-treatment did not impact worms’ overall translational movement on the chemotaxis plates (Fig 4C), indicating that NAM causes a specific defect in chemotaxis rather than a general depression in movement ability.

**Figure 4.**
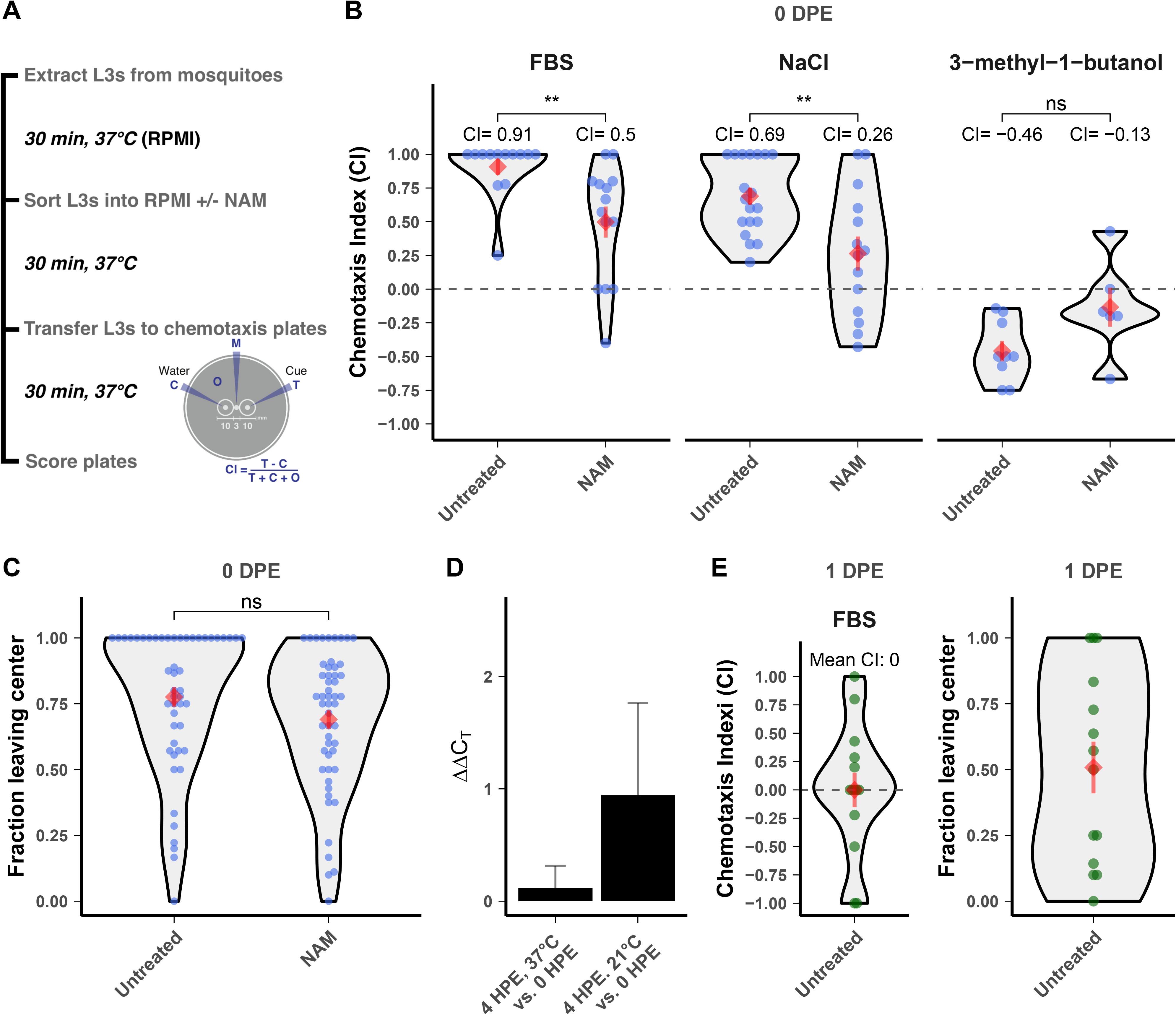
Treatment with a TRVP agonist dysregulates chemotaxis of *Brugia pahangi* infective larvae. **(A)** L3 stage parasites were extracted from mosquitoes and subjected to chemotaxis assays with or without 250 μM nicotinamide (NAM) treatment. Chemotaxis assays are performed by adding L3s to the center of a 0.8% agarose plate (M), either test cue (T) or water (C) to the opposite sides of the plate, and placed at 37°C for 30 minutes. Plates are scored after incubation and the chemotaxis index (CI) is calculated, **(B)** NAM dysregulates attraction of freshly extracted L3s to serum and NaCI, but has no effect on aversion to 3-methyl-1-butanol. **(C)** NAM has no effect on gross translational movement of freshly extracted L3s. **(D)** *Bpa-osm-9* expression is unchanged by *in vitro* culture at physiological or room temperature 4 hours post-extraction (HPE). **(E)** L3s cultured for 1-day post-extraction (DPE) do not show chemotaxis toward serum and have reduced motility on the chemotaxis plate when compared to untreated freshly extracted parasites (p = 0.028, t-test). Data for A-C represent the combined results of three independent biological replicates, except for the experiments with 3-methyl-1-butanol, which included two replicates (cohorts of mosquito infections). Data for D represents the results of two biological replicates. Each point represents a single chemotaxis plate with 8-10 L3s. Red diamonds and bars indicate the mean and standard error of the mean. Comparisons of means were performed using t-tests (**: p <= 0.01).

To ensure that the *Bpa-osm-9*, the putative target of NAM, was expressed during the performance window of our assay and that expression was not altered by the ambient temperatures that L3s experience during assay preparation, we measured the relative expression of *Bpa-osm-9* in L3s immediately after extraction from mosquitoes, and after 4 hours of *in vitro* culture at human body temperature (37°C) or ambient temperature (~21°C). The relative expression of *Bpa-osm-9* was unchanged over this time frame at either temperature (p = 0.4215, Fig 4D).

Although the expression of *Bpa-osm-9* does not drastically change following extraction (Fig 4D), parasites maintained overnight under standard culture conditions do not show a chemotactic response to serum, even with pre-assay incubation in serum-free media, and show a reduced motility on the chemotaxis plate when compared to untreated freshly extracted parasites (Fig 4E). Although it is possible that the specific unknown chemoreceptors involved in serum response are downregulated by this time point, it is more likely that artificial culture conditions have effects on parasite health that compromise chemotactic potential. These results highlight the importance of using freshly extracted L3 larvae in these assays.

### *Treatment with a nematode TRPV agonist alters infective-stage* Brugia *larvae thermosensory response*

L3 stage larvae that have departed the intermediate mosquito host are challenged with stark temperature shifts from the ambient temperature in the mosquito, to warmer temperatures on the definitive host’s skin ~24-34°C (57), to an even warmer still host core temperature of 37°C. During *in vitro* culture, healthy L3 worms elongate and vigorously thrash in media (Fig 5B, bottom), but thrashing will transition to coiling and reduced motility as culture media cools (Fig 5B, top). The coiling and reduction in motility caused by cooling is reversed after returning the parasites to 37°C (S4 Figure), indicating that the phenotype is not a result of general sickness or tissue damage. In the course of performing L3 chemotaxis experiments with NAM, we noticed that treated L3s had a reduced coiling response. To confirm this effect, we performed dose-response experiments and video-recorded parasites exactly 20 minutes after transfer from 37°C to room temperature, at the point where untreated parasites tightly coil. Both blinded manual scoring and a bespoke computer imaging analysis of larval coiling reveals that NAM inhibits this thermosensory response in a dose-dependent manner after 24 and 48 hours (Fig 5C-D). These data suggest that *Brugia* OSM-9 plays a polymodal sensory role in L3 stage parasites, potentially mediating both chemosensory and thermosensory responses.

**Figure 5.**
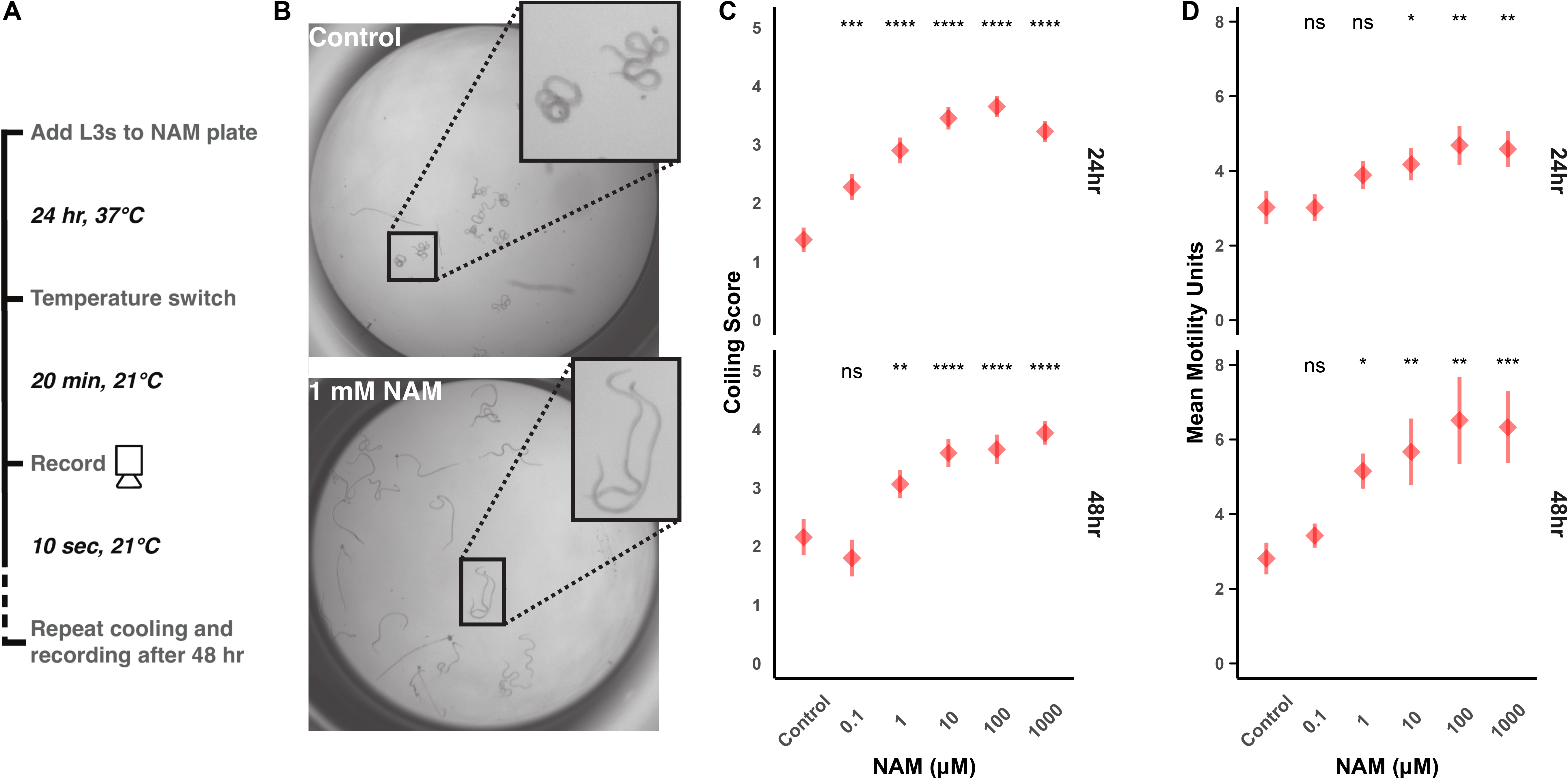
Treatment with a TRPV agonist impairs the temperature-shift coiling response in *Brugia pahangi* infective larvae. **(A)** L3 stage parasites were extracted from mosquitoes and treated with NAM, subjected to a temperature shift, and analyzed for cooling-induced coiling behaviors. **(B)** Representative images of untreated (control) individuals displaying the coiled phenotype and 1 mM NAM exposed individuals with uncoiled thrashing. **(C)** Blinded coiling score given to each treatment after 24 hours and 48 hours post-treatment (higher score indicates less coiling). **(D)** Mean motility calculated by an optical flow algorithm. Red diamonds and bars indicate the mean and standard error of the mean from three biological replicates, each composed of >3 technical replicates scored by three different researchers. Comparisons of means were performed using one-sided t-tests. (*: p <= 0.05; **: p <= 0.01, ***: p <= 0.001; ****: p <= 0.0001)

### Pretreatment of microfilariae with NAM reduces L3 burden in infected mosquitoes and alters tissue distribution

Assays with extracted L3s indicated that Bpa-OSM-9 may be important for *in vitro* chemoattraction to salt and serum. We hypothesized that NAM could dysregulate intra-mosquito chemotaxis of larval stages *in vivo*. To establish an assay to test this hypothesis, we first investigated whether NAM had any effect on mosquito blood-feeding dynamics. When NAM was added to blood and provided to mosquitoes to feed *ad libitum*, 0.1 μM to 5 mM NAM acted as a phagostimulant, causing a dose-dependent increase in the proportion of mosquitoes that had fed after 30 minutes. However, feeding proportions began to decrease at concentrations greater than 5 mM, and 250 mM NAM caused complete repulsion to the blood (Fig 6A). We chose 5 mM and 25 mM as initial treatment concentrations for mf, and with replication we found that 5 mM NAM caused a significant increase in the proportion of mosquitoes that fed, while 25 mM caused no significant difference from unsupplemented blood (Fig 6B). To ensure that the increase in proportion of fed mosquitoes was not correlated to an increased blood meal size, we measured distended abdomens of mosquitoes after feeding on control blood or blood supplemented with 5 mM or 25 mM NAM (S5 Figure). Mosquito abdomen sizes were unchanged by NAM supplementation, assuring that altered parasite burdens after feeding would not be a function of altered numbers of ingested mf (Fig 6C).

**Figure 6.**
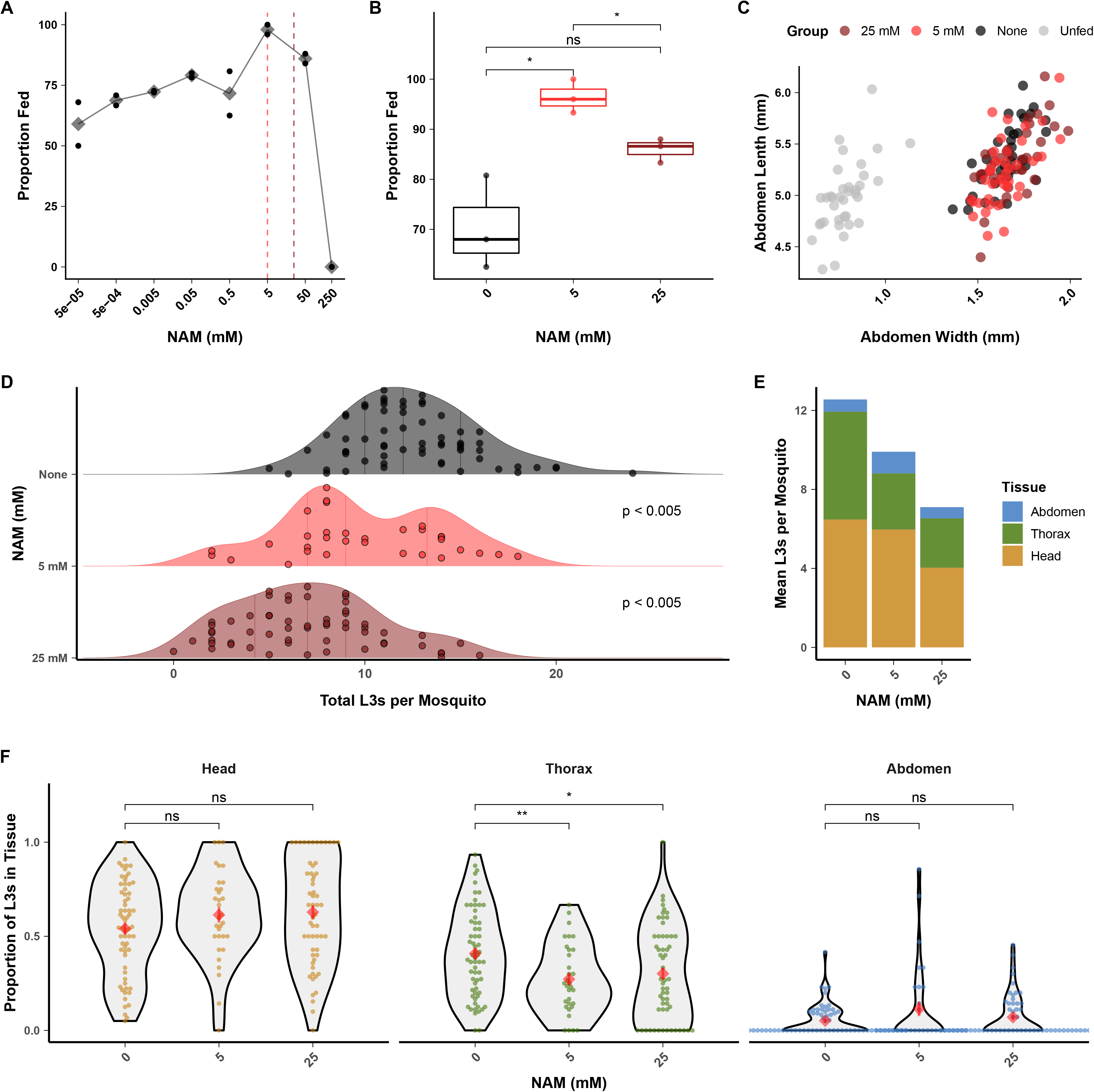
Treatment with a TRPV agonist reduces the ability of microfilariae to establish infection in mosquitoes. **(A)** Nicotinamide (NAM) added to blood at up to 50 mM increases the proportion of blood-fed mosquitoes when allowed to feed to repletion, but reduces mosquito blood-feeding at concentrations greater than 5 mM. Black points represent technical replicates and grey diamonds represent the mean. **(B)** Replication of blood-feeding experiments with 5 mM and 25 mM showed a significant increase in the proportion of blood-fed mosquitoes when blood was supplemented with 5 mM NAM, but no difference when supplemented with 25 mM. These concentrations were used for subsequent parasite treatment. Points represent the values from three independent biological replicates (cohorts of mosquitoes). **(C)** Blood supplemented with 5 or 25 mM NAM does not alter the size of distended mosquito abdomens after blood-feeding, indicating an unaltered size of blood meal. Points represents the measured abdomens of individual mosquitoes from a single blood-feeding experiment. **(D-E)** Pre-treatment of *B. pahangi* microfilaria (mf) with NAM prior to mosquito infection causes a dose-dependent reduction in the number of L3s recovered per mosquito. P-values were calculated with Tukey’s post-hoc tests and adjusted for multiple comparisons. **(F)** Reduction in L3 recovery was due to a decrease in larval parasites in the mosquito thorax (i.e. the flight muscles, the migratory destination for mf and site of development for L1, L2, and early-L3 stage parasites). Data from D-F represents the combined results of three independent biological replicates (cohorts of mosquito infections); each point represents the parasites recovered from an individual mosquito. Red diamonds and bars indicate the mean and standard error of the mean. Comparisons of means in B and F were performed using t-tests. (*: p <= 0.05; **: p<= 0.01)

We next supplemented microfilaremic blood with 5 mM and 25 mM NAM and tested for altered infectivity and intra-mosquito tissue distribution of larvae. Pretreatment with NAM of *B. pahangi* mf caused a significant, dose-dependent reduction in parasite burden at 14 DPI (Fig 6D-E, 5 mM = 21% reduction, 25 mM = 43% reduction) and a significant decrease in the proportion of L3s recovered in the thorax of infected mosquitoes (Fig 6F). The proportion of L3s recovered in the thorax was not correlated to total L3s recovered per mosquito (S6 Figure), suggesting that changes in larval infectivity were not due to differences in blood meal size, but instead the specific action of NAM upon the parasite. Thus, we postulate that NAM inhibits the initial migration of mf from the blood bolus, but that once across (and presumably relieved of NAM exposure in the midgut), developed L3 larval parasites are able to migrate to the head at the same proportion as untreated controls.

### Brugia osm-9 and tax-4 RNAi inhibits chemoattraction of infective-stage larvae toward host-associated cues

NAM is an agonist of the *C. elegans* TRPV heteromer OSM-9/OCR-4 and *Drosophila* orthologs Nanchung/lnactive when expressed in *Xenopus* oocytes, but not of either *C. elegans* subunit alone (56). Given conservation of NAM-receptor interactions across these phyla, we expect *Brugia* OSM-9 orthologs to also respond to NAM. However, the pharmacology and subunit interactions of *Brugia* OSM-9 may differ (e.g., filarial parasites do not have a homolog of *ocr-4* (Fig 3A)), and NAM is an endogenous metabolite in *C. elegans* that has pleiotropic effects (56,58–62). This compelled us to use a genetic approach to more directly test whether *Brugia* OSM-9 and TAX-4 are involved in parasite chemotaxis behavior.

We carried out intra-mosquito RNA interference (RNAi) (63) of both *Bpa-osm-9* and *Bpa-tax-4* in larval stages and measured the effects on L3 *in vitro* chemotaxis. Infected mosquitoes were injected with dsRNA targeting transcripts of interest at 9 DPI, corresponding to the expected timeline of L2 to L3 transition in the thoracic musculature (8) (Fig 7A). We attempted to confirm knockdown of target transcripts with qPCR, but the low target abundance of *Bpa-osm-9* and *Bpa-tax-4* relative to housekeeping genes, coupled with limited recovery of RNA from a small number of parasites prevented reliable amplification. Targeting either *Bpa-osm-9* or *Bpa-tax-4* using the *‘in squito’* RNAi protocol resulted in the inhibition of *B. pahangi* L3 *in vitro* chemotaxis towards serum at 14 DPI (Fig 7B), while injection of non-specific (*lacZ*) dsRNA had no effect on chemotaxis (control chemotaxis index (CI): 0.83, *Bpa-osm-9(RNAi*) CI: 0.21, *Bpa-tax-4(RNAi*) CI: 0.12). dsRNA treatment did not have any effect on general parasite motility on the assay plate (Fig 6C). To our knowledge, this is the first time that either *tax-4* or *osm-9* have been shown to have a specific function in chemosensation in a parasitic nematode.

**Figure 7.**
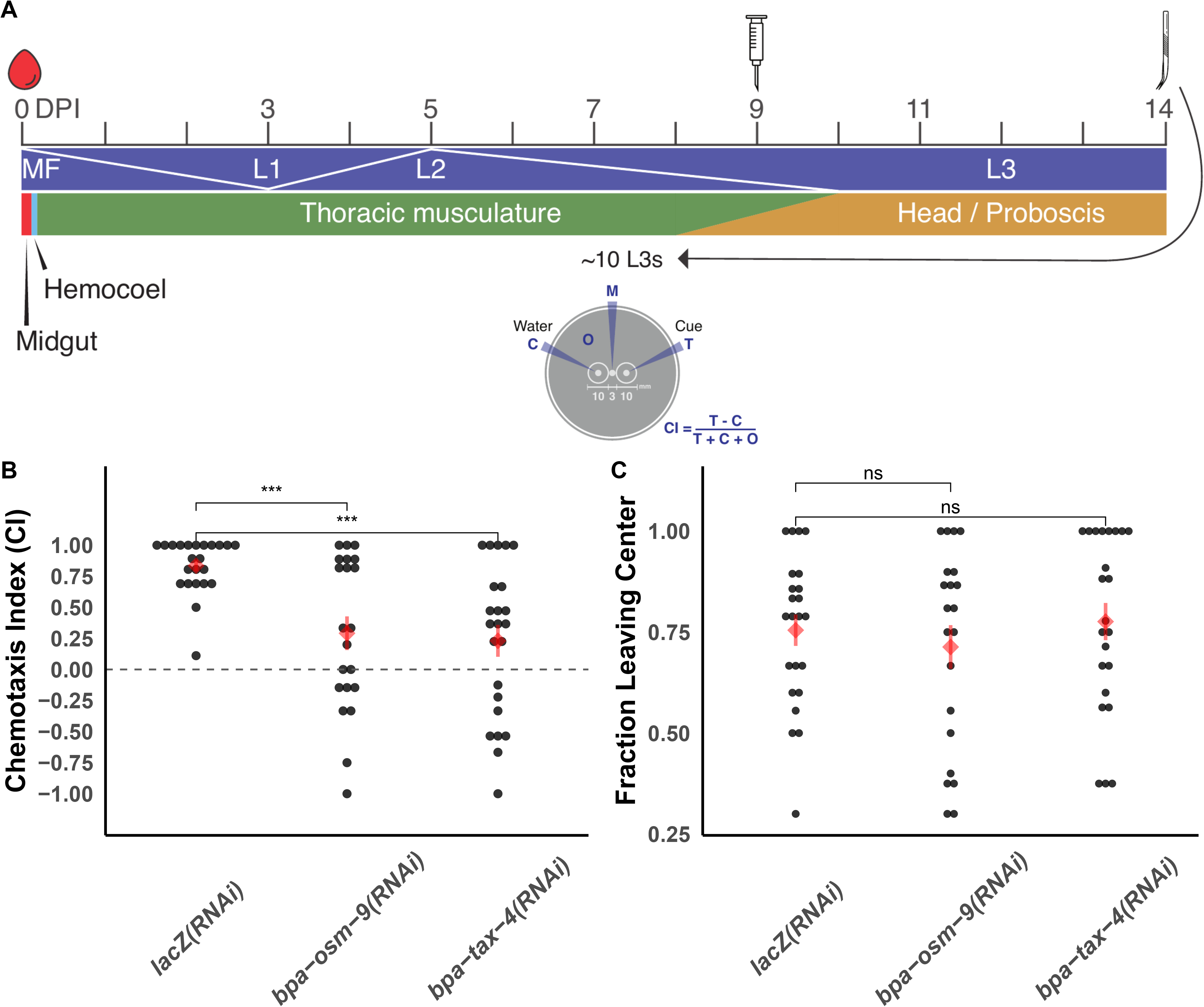
dsRNA treatment of chemosensory pathway receptors causes defective chemotaxis of *Brugia pahangi* infective larvae. **(A)** Injection of 250 ng dsRNA into *B. pahangi*-infected *Ae. aegypti* LVP was performed 9 DPI, and L3 parasites were recovered via dissection at 14 DPI. Recovered parasites were immediately used in chemotaxis experiments. Chemotaxis assays are performed by adding L3s to the center of a 0.8% agarose plate (M), either test cue (T) or water (C) to the opposite sides of the plate, and placed at 37°C for 30 minutes. Plates are scored after incubation and the chemotaxis index (CI) is calculated, Intra-mosquito developmental dynamics adapted from (8). **(B)** dsRNA treatment of *Bpa-osm-9* or *Bpa-tax-4* resulted in a reduced ability of L3s to migrate to serum. Control parasites were recovered from mosquitoes injected with *lacZ* dsRNA. dsRNA exposure does not inhibit general translational motility on the chemotaxis plate. Data represents the combined results of three independent biological replicates (cohorts of mosquito infections); each point represents the chemotaxis index of an individual plate. Red diamonds and bars indicate the mean and standard error of the mean. Comparisons of means were performed using t-tests. (***: p <= 0.001; ****: p <= 0.0001)

### *Homologous TRP and CNG channels from* Brugia malayi *partially rescue sensory defects in* C. elegans

To further explore the sensory functions of *Brugia osm-9* and *tax-4, we* tested whether these genes could rescue behavioral defects in *C. elegans* strains with loss-of-function mutations in endogenous *osm-9* or *tax-4*. While the sequenced genome of *B. malayi* is near completion, many of the gene models remain fragmented and unconfirmed, so we performed low-coverage isoform sequencing with long-read RNA-seq on *B. malayi* adult males and females. This led to the successful capture of *Bma-osm-9* full-length transcripts, but failed to capture *Bma-tax-4*. Using these data and the predicted gene model of *Bma-tax-4*, we cloned these genes and for functional expression in *C. elegans We* also cloned the *Bma-ocr-1/2*-like gene that has the highest predicted amino acid identity to OCR-2 (*Bm5691*, or *Bma-ocr-1/2a*), which functions with OSM-9 in *C. elegans* to enable a range of sensory behaviors.

The *Bma-osm-9* clone and the two full length isoforms captured by long-read sequencing included a 41 bp insertion that corresponded to a missing splice-acceptor site at intron 17 that was not reflected in the original gene prediction (S7 Figure). This insertion caused a frame-shift in the predicted amino acid sequence that made the resulting sequence more similar to the *Cel-osm-9* sequence than the original prediction (S8 Figure). The consensus *Bma-tax-4* transcript we cloned had two differences from the predicted model(38), a synonymous 694T>C that was found in 4 out of 7 sequenced clones, and a 21 bp deletion that was found in all clones and corresponds to a mispredicted splice-donor site at intron 2 (S9 Figure). The consensus *Bma-ocr-1/2a* sequence was a perfect match to the predicted gene model. We used these clones to carry out a range of sensory assays to test for rescue of *C. elegans* chemotaxis defects by the *B. malayi* homologs.

These genes were initially expressed in corresponding loss-of-function *C. elegans* backgrounds (*osm-9(ky10*) and *tax-4(p678)*) using *Cel-osm-9* and *Cel-tax-4* promoter regions and the commonly used *unc-54* 3’ UTR (28,30). These transgenes did not rescue defects in chemotaxis to diacetyl, nose-touch reversal, or benzaldehyde avoidance, and neither did co-expression of *Bma-osm-9* and *Bma-ocr-1/2a* (Fig 8A-C). Inability to rescue was not due to a lack of mRNA expression (S10 Figure). We hypothesized that although the parasite transgenes were expressed generally, they may not have been expressed in the neurons required for chemotaxis (AWA) and avoidance (ASH) due to a lack of *cis*-regulatory elements that control expression in these neurons (30). We subsequently replaced the *unc-54* 3’ UTR with 3.2 kb of the *Cel-osm-9* downstream region and found that this modification allowed for partial rescue of the avoidance defects but not of chemotaxis defects. Self-rescue with *C. elegans osm-9* also did not rescue the chemotaxis defect, suggesting that there are addition regulatory elements that enable expression in AWA. Likewise, *Bma-tax-4* showed partial rescue of chemotaxis to isoamyl alcohol, which is controlled by the AWC neuron, even without *Cel-tax-4* 3’ *cis*-regulatory elements (Fig 9). These results show that *Brugia* OSM-9 and TAX-4 are able to partially rescue *C. elegans* sensory defects by functioning as channel subunits in a free-living nematode cell context, suggesting functional conservation across diverged nematode species, while the failure to completely rescue may point to some amount of subfunctionalization.

**Figure 8.**
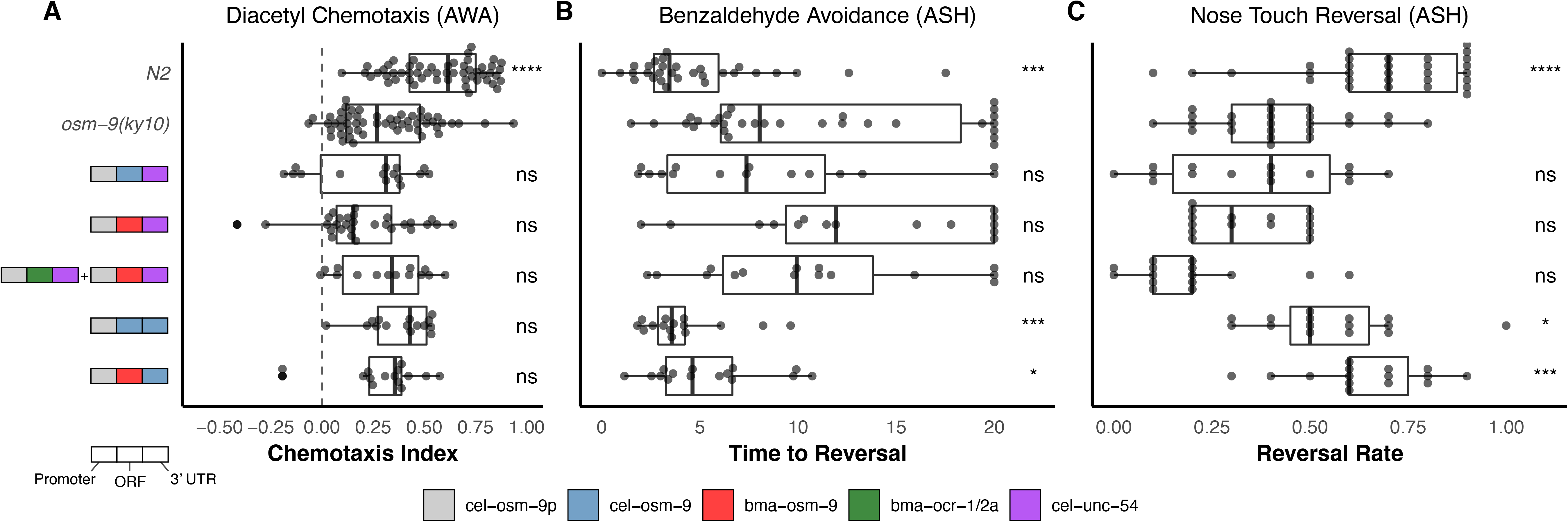
Heterologous expression of *B. malayi osm-9* partially rescues loss-of-function sensory defects in *C. elegans*. *Bma-osm-9* and *Bma-ocr-1/2a* were cloned and expressed under the control of the endogenous *Cel-osm-9* promoter (28,30). The *Cel-osm-9* open-reading frame was used as a positive control **(A)** Chemotaxis defects to diacetyl (OSM-9 functioning in AWA) was not rescued by any of the constructs, while avoidance defects (OSM-9 functioning in ASH) to **(B)** concentrated benzaldehyde and **(C)** mechanical nose touch were partially rescued by the positive control and by *Bma-osm-9* with *Cel-osm-9* 3’ UTR. Data represents the combined results of at least three independent biological replicates, each consisting of 5 technical replicates. Each point represents the chemotaxis index of an individual plate or worm, depending on the assay. Comparisons to the loss-of-function strain were performed using t-tests. (****: p <= 0.0001, ***: p <= 0.001, **: p <= 0.01, *: p <= 0.05)

**Figure 9.**
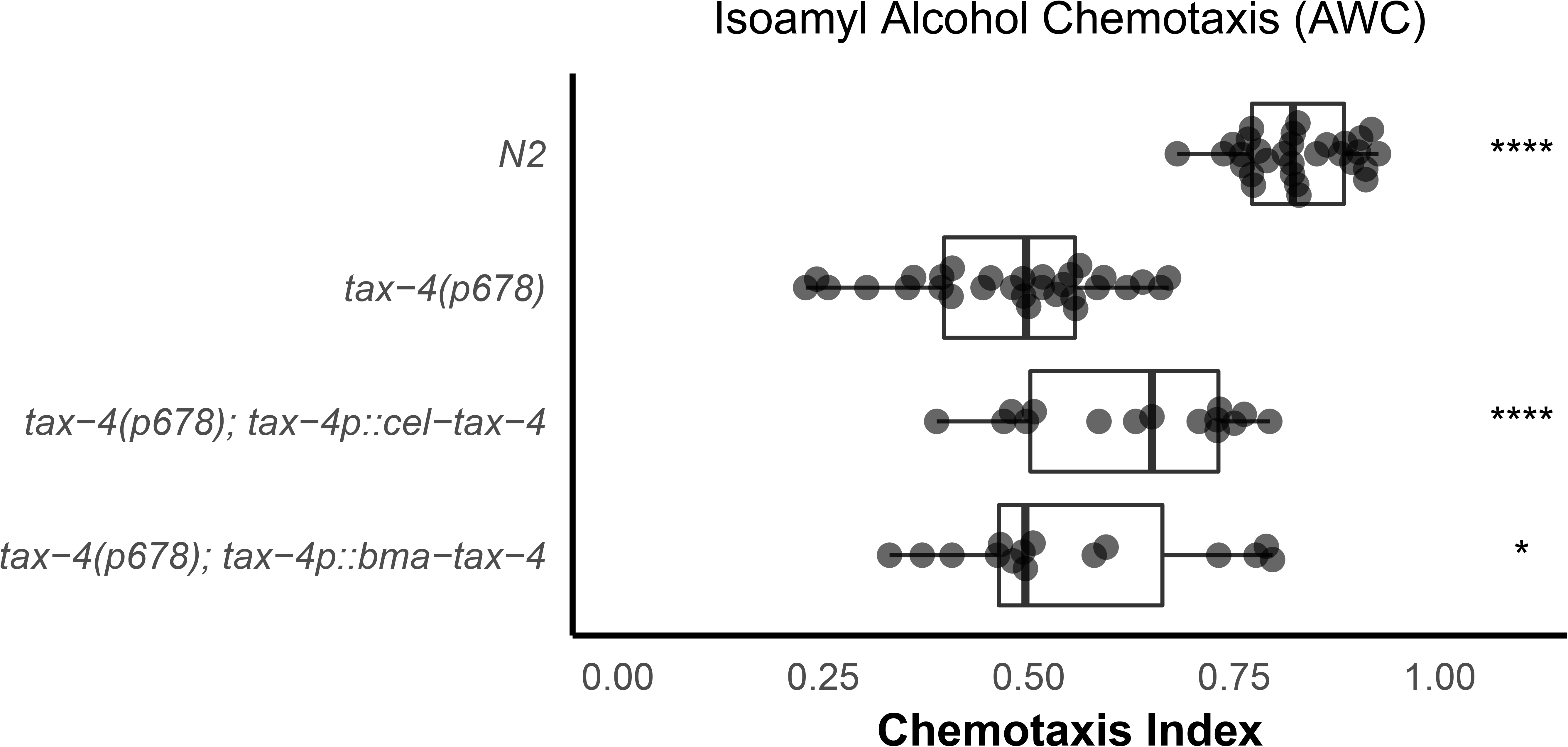
Heterologous expression of *B. malayi tax-4* partially rescues loss-of-function chemotaxis defects in *C. elegans*. *Bma-tax-4* was cloned and expressed under the control of the endogenous *Cel-tax-4* promoter. The *Cel-tax-4* open-reading frame was used as a positive control. Both constructs partially rescued the chemotaxis defect to isoamyl alcohol (TAX-4 functioning in AWC). Comparisons to the loss-of-function strain were performed using t-tests. (****: p <= 0.0001, *: p <= 0.05)

## Discussion

Filarial worms continue to pose a significant threat to human and animal health. The ability for filarial worms to move through and between hosts relies on their ability to sense their environment, evidenced by the diversity of genus-specific niches occupied by different filariae when they migrate within shared arthropod or vertebrate hosts (e.g. *Brugia* larvae migrate to the thoracic musculature of *Ae. aegypti*, while *Dirofilaria* migrate to the Malphigian tubules). Despite the importance of sensory behaviors in the evolution and persistence of parasitism, little is known of the receptors and pathways that control such behaviors in parasitic nematodes. Identifying mediators of sensory-associated behaviors can aid our understanding of disease transmission and pathogenesis, and may also provide new targets for therapeutic intervention.

We have shown that filarial worms have a greatly reduced and divergent set of chemoreceptors as compared to *C. elegans* and the rest of the Nematoda, but that they retain much of the core downstream chemosensory signaling pathway. Filarial parasites exhibit stage, tissue, and sex-specific chemoreceptor patterns that likely correspond to the different vector and host environments encountered throughout the life cycle. We expect that these patterns correspond to landmark migration events, including the migration of larvae within the mosquito host, the transmission of infective larvae (L3s) to the definitive host, the early migration of larvae in the definitive host, and the potential mate-seeking behaviors of dioecious adults.

Historically, it has been difficult to identify endogenous ligands for nematode chemoreceptors. In *C. elegans*, only a small fraction of chemoreceptors have been linked to activating molecules, some of which are unlikely to be the natural ligand (49,64–73). This is partly a function of the large number of chemoreceptors (>1,400) and the large space of potentially relevant terrestrial cues in free-living clade V nematodes. The smaller complement of chemoreceptors in clade IIIc filarial parasites combined with the stark temporal receptor expression patterns overlayed with possible host-derived molecules present at intra-host sites of parasite migration may facilitate a comparatively easier path to chemoreceptor deorphanization. Efforts are underway to develop new heterologous expression platforms for deorphanization that may be more amenable to the expression of nematode GPCRs, which have often been recalcitrant to expression in single-cell mammalian or yeast systems.

As in *C. elegans*, TRP and CNG channels in filarial worms likely function downstream of chemoreceptors expressed on the cilia of sensory amphids. We show that a TRPV chemical agonist inhibits *in vitro* chemotaxis of infective *Brugia* larvae towards host-associated cues and compromises the ability of microfilariae to establish mosquito infections. RNAi experiments implicate both *Brugia* OSM-9 and TAX-4 as necessary for larval chemotaxis to serum. In *C. elegans*, OSM-9 and TAX-4 function as primary sensory transducers in distinct sensory cells that can respond to distinct chemicals. It is interesting that both OSM-9 and TAX-4 mediate responses to serum in *B. pahangi*. FBS is a complex heterogeneous mixture of macromolecules, amino acids, and ions, and it is conceivable that different components of this mixture activate distinct chemoreceptors, chemosensory neurons, and downstream pathways in *Brugia*. Little is known about the neuronal architecture of filarial worms, and it is also possible that OSM-9 and TAX-4 are coexpressed and have homologous function in the same cells. A map of the filarial worm connectome and the ability to produce transcriptional reporters would help resolve this question, and will be hastened by technologies to more easily dissect filarial worm neuroanatomy and neurogenetics (74). Furthermore, fractionation of serum into pure chemicals for chemotaxis experiments could illuminate whether multiple cells are involved in *Brugia* chemotactic responses to serum or other crude preparations.

CNG and TRP channels are polymodal in *C. elegans* and function in sensory neurons responsible for aerosensation, mechanosensation, chemosensation, noxious avoidance, and thermosensation, among others (28–30,75–77). Our data suggest a similar polymodal deployment of these channels in mosquito-borne filarial worms, though the pattern of neuronal expression may differ. While OSM-9 in *C. elegans* is involved in noxious heat avoidance in the nociceptive ASH neuron (76), it is TAX-4 and TAX-2 that function in the sensation of precise of thermal gradients via AFD. Whether filarial worms have cooperative thermal sensory programs is unknown, but given the tight range of temperatures experienced by these parasites (from ambient temperatures while in mosquitoes to physiological temperatures in mammalian hosts) and the unlikelihood of experiencing or being able to avoid noxious heat or cold, the sensory program in filarial worms is likely more simple than their free-living counterparts. Our data suggest that OSM-9 is involved in this program, but CNGs like TAX-4/TAX-2 cannot be ruled out.

Questions remain as to the stoichiometry and subunit interactions of TRP and CNG channels that function in *Brugia* chemosensation. Both *Bma-osm-9* and *Bma-tax-4* were able to rescue sensory defects in *C. elegans* without co-expression of putative subunits (e.g. *ocr-2* and *tax-2*). so it is possible that the parasite subunits were able to form heteromeric channels with their free-living counterparts, or that the parasite subunits were able to form homomeric channels. Regardless, the clade IIIc loss of CNGs involved in olfactory plasticity (*cng-1, cng-3*) and TRPs that are expressed in the mechanosensory labial QLQ neurons of *C. elegans (ocr-4, trpa-1*) demonstrate imperfect conservation of all sensory modalities and pathways between *C. elegans* and filarial worms, and it is possible that filarial worm TRP and CNG channels have evolved subunit interactions or primary functions that are not conserved in *C. elegans* (31,53,78,79).

Deeper knowledge of chemotaxis has been achieved in clade IV and V nematodes that are more amenable than filarial worms to *in vitro* culture and manipulation (15,80). Continued development of genetic tools (63,74,81) and sensory assays would help to further elucidate the molecular basis for sensory behaviors in clade III LF parasites. The *in vitro* chemotactic capacity of filarial worm L3s is transitory, and adults, which are healthy in culture much longer than larval parasites, could offer an additional platform for sensory pathway dissection. Exploration of other sensory modalities such as thermosensation and mechanosensation are essential to develop a more thorough model of how filarial worms integrate sensory data in order to successfully invade, infect, and migrate within the host.

## Materials and Methods

### Protocol and data availability

All comparative genomics, phylogenetics, data analysis, and data visualization pipelines are publicly available at https://github.com/zamanianlab/BrugiaChemo-ms. The optical flow algorithm for motility analysis is available at https://github.com/zamanianlab/BrugiaMotilityAnalysis. Short-read and long-read sequencing data has been deposited into NIH BioProjects <u>PRJNA548881</u> and <u>PRJNA548902</u>, respectively.

### Parasite and mosquito maintenance

Microfilariae (mf), L3, and adult stage FR3 strains of *Brugia malayi* and *Brugia pahangi* from the NIH/NIAID Filariasis Research Reagent Resource Center (FR3) (82) were maintained in RPMI 1640 culture media (Sigma-Aldrich, St. Louis, MO) with penicillin /streptomycin (0.1 mg/mL, Gibco, Gaithersburg, MD) and FBS (10% v/v, Gibco) unless otherwise stated. For local production of L3 stage *B. pahangi*, mf were incubated in defibrinated sheep’s blood (Hemostat, Dixon, CA) at a density of 120-160 mf / 20 μl at 37°C in a membrane feeder(83). mf were exposed to groups of 250 adult female *Aedes aegypti* Liverpool strain (LVP) 1-3 day post-emergence starved 1 day prior to feeding. Infected mosquitoes were maintained in double housed cages in a Percival Scientific incubator (Perry, lA) model I-36NL incubator (26°C, 85% humidity, 12:12 hr light:dark cycle) and provided 10% sucrose throughout.

At 14 days post-infection (DPI) L3 stage parasites were extracted into warm *Aedes* saline (84) or RPMI 1640 (Sigma-Aldrich) via micro-dissection of cold anesthetized mosquitoes or bulk isolation as previously described(9). The prevalence and locality of L3s was determined by separating head, thorax, and abdominal tissues during dissection with all L3s counted per mosquito. *B. malayi* adults used for RNA-seq were received from the FR3 and were immediately washed and placed in new media. Adult parasites were allowed to equilibrate at 37°C for 24 hours before any further experimentation.

### Caenorhabditis elegans *strains*

*C. elegans* strains were maintained at 20°C on NGM plates seeded with *E. coli* strain OP50 and routinely picked to fresh plates at the L4 stage. Transgenic strains were created as described(85) by microinjecting 50 ng/uL of parasite transgene, combined with either *unc-122p::GFP* or *myo-2p::GFP* as co-injection markers and an empty vector to achieve a final concentration of 100 ng/uL. Three independently derived lines were created for each transgenic strain and maintained separately. Genotypes used include: *osm-9(ky10*) IV, *tax-4(p678*) III, ZAM13: *osm-9(ky10*) IV, *mazEx13[osm-9p::Bma-osm-9::unc-54* 3’UTR; *unc-122p::GFP*], ZAM14: *tax-4(p678*) III, *mazEx14[tax-4p::Bma-tax-4::unc-54* 3’UTR; *myo-2p::GFP*], ZAM17: *osm-9(ky10*) IV, *mazEx13[osm-9p::Bma-osm-9::unc-54* 3’UTR; *unc-122p::GFP*], *mazEx17[osm-9p::Bma-ocr-1/2a::unc-54* 3’UTR; *myo-2p::GFP*], ZAM18: *osm-9(ky10*) IV, *mazEx18[osm-9p::Cel-osm-9::unc-54* 3’UTR; *unc-122p::GFP*] ZAM21: *tax-4(p678*) III, *mazEx19[tax-4p::Cel-tax-4::unc-54* 3’ UTR; *myo-2p::GFP*], ZAM22: *osm-9(ky10*) IV, *mazEx20[osm-9p::Bma-osm-9::osm-9* 3’UTR; *myo-2p::GFP*,] ZAM24: *osm-9(ky10*) IV, *mazEx21[osm-9p::Cel-osm-9::osm-9* 3’UTR; *myo-2p::GFP*].

mRNA expression of some B. malayi genes in transgenic C. elegans was confirmed with qPCR (S10 Figure; primer sequences are included in S4 File). A 20 μL qPCR reaction was optimized using 2X PowerUp SYBR Green MasterMix (Thermo Fisher Scientific) and 10 ng RNA isolated from mixed populations from 2-3 chunked plates as described(86) converted to cDNA with SuperScript III using an equal amount of random hexamer and oligo(dT) primers. Reactions were run in duplicate on a StepOnePlus real-time PCR system (Applied Biosystems, Waltham, Massachusetts). C_T_ values were calculated with the system’s automatic threshold, and relative expression was calculated with the ΔΔC_T_ method(87). Parasite transcripts in negative controls (knock-out C. elegans strains) were undetected in all cases and C_T_ was set to 40 to calculate a ΔΔC_T_ for data visualization.

### Comparative genomics

#### Chemosensory GPCRs

The chemoreceptor mining and annotation strategy is charted in S1 Figure. Briefly, all filarial worm predicted proteomes in WormBase ParaSite version 9(38) and a selected list of high-quality genomes that included representatives from the four major nematode clades (39 total species, S1 Table) were searched (hmmsearch(88)) against a database of profile hidden Markov models (HMMs) curated by Pfam, consisting of primary metazoan GPCR families and the nematode chemoreceptor families(89). Predicted proteins were filtered such that each that had a best-hit to a nematode chemoreceptor HMM was retained. Surviving predicted proteins were then used in a reciprocal search (hmmsearch) against the entire Pfam HMM database. Predicted proteins that had a best-hit to a nematode chemoreceptor HMM were retained. Surviving predicted proteins were then searched (blastp(90)) against the *C. elegans* predicted proteome (N2, WBPS9), and predicted proteins that had a best-hit to a *C. elegans* chemoreceptor (S4 Data) were retained (C. *elegans* chemoreceptors were obtained by downloading all named chemoreceptors from WormBase and their paralogues via the WormBase ParaSite API).

#### TRP and CNG receptors

Predicted protein sequences of annotated TRP and CNG channels from *C. elegans were* downloaded from WormBase(91) and used as seeds in blastp searches against all predicted proteomes included in S1 Table. Hits with an E-value < 0.01 were reciprocally searched against the *C. elegans* predicted proteome, and any hit that wasn’t most similar to a *C. elegans* TRP or CNG channel was removed. Because there were clade and species-specific gene losses in the CNG family, *C. elegans* seeds were also used in a tblastn search against parasite genomes to account for missing gene models and possible errors in gene predictions.

### Phylogenetics

#### Chemosensory GPCRs

Predicted protein sequences belonging to *C. elegans* chemoreceptor families (39) were aligned by family with MAFFT(92). The resulting family profile HMMs were sequentially aligned with MUSCLE(93) to create a master *C. elegans* chemoreceptor alignment. Predicted chemoreceptors from 19 selected species underwent transmembrane domain prediction with HMMTOP(94), and only those that contained exactly 7 predicted TMs were aligned to the master alignment. This final multiple sequence alignment was trimmed with trimAI(95) such that columns with greater than 30% gaps were removed, and sequences that did not have at least 70% of residues that aligned to columns supported by 70% of the sequences were removed.

The trimmed, filtered alignment was subjected to maximum-likelihood phylogenetic inference with IQ-TREE(96) and ModelFinder(97) with ultrafast bootstrapping(98), using the VT substitution matrix(99) with empirical base frequencies and a free-rate substitution model(100,101) with 10 categories. Bootstrap values from 1000 replicates were drawn as nodal support onto the maximum-likelihood tree.

#### TRP and CNG receptors

Putative TRP sequences underwent TM prediction, and any sequences with ≥1 predicted TMs were retained. TRP and CNG sequences were separately aligned, trimmed such that columns with greater than 25% gaps were removed, and CNG sequences that did not have at least 70% of residues that aligned columns supported by 70% of the sequences were removed. For both datasets, fragments with large gaps or putative isoforms were manually removed. Alignments were subjected to Bayesian phylogenetic inference with MrBayes(102). The MCMC chain was run for 10,000,000 generations, with a sample taken every 1000 generations. Eight separate chains were run, with two hot chains and the temperature set to 0.05. Consensus trees were drawn using the 50% majority rule, with all compatible groups added, and posterior probabilities were drawn as nodal support. All trees were annotated with ggtree(103).

### Brugia malayi *transcriptomic analyses*

#### Anterior and posterior *B. malayi* transcripts

One millimeter of the anterior and posterior ends of 19 adult male and 18 adult female *B. malayi* were cut from live parasites and immediately transferred to Trizol (Ambion, Waltham, MA). Tissue in Trizol was homogenized with a plastic pestle, and RNA was extracted with Direct-zol RNA miniprep kit (Zymo, Irvine, California) according to the manufacturer’s instructions and was eluted in RNase-free water. RNA was DNase-treated on the column, and the quality of purified RNA was assessed with Qubit (Thermo Fisher Scientific, Waltham, MA) and Bioanalyzer Pico chip (Agilent, Santa Clara, CA). RNA was rRNA depleted with Ribo-Zero ScriptSeq Complete Gold (Blood) (Illumina, San Diego, CA) and sequencing libraries were constructed using the TruSeq Stranded Total RNA kit (Illumina). All samples were sequenced at the UW-Madison Biotechnology Center with an Illumina HiSeq 2500 with a single-end 100 bp read setting. Reads were adapter and quality trimmed using Trimmomatic(104). HiSAT2(105) and StringTie(106) were used to align reads to the *B. malayi* reference genome (WormBase ParaSite(38), release 12 version 4) and to produce TPM (transcripts per million) counts for annotated genes. The RNA-seq pipeline was implemented using Nextflow(107) and is publicly available (https://github.com/zamanianlab/Bmalayi_HTRNAseg-nf). Custom R scripts were used for profiling, hierarchical clustering, and visualization of putative chemosensory gene expression across anterior and posterior samples.

#### Stage-specific expression of *B. malayi* chemosensory genes

Public stage-specific RNA-seq data(52) was acquired from NCBI SRA. Reads were aligned to version 4 of the *B. malayi* genome, downloaded from version 12 of WormBase ParaSite(38). Reads were aligned with HISAT2(105) and StringTie(106). Custom R scripts were used for profiling, hierarchical clustering, and visualization of putative chemosensory gene expression across life stages. Heatmaps of life stage expression were drawn according to chromosomal location. The RNA-seq pipeline was implemented using Nextflow(107) and is publicly available (https://github.com/zamanianlab/BmalayiRNAseg-nf). Locus information was extracted from GTF files from WormBase ParaSite(38) and plots were generated with Circos(108).

### Brugia *chemotaxis assays*

All chemotaxis assays were performed immediately after extraction of L3s from local infections and following previously published protocols(23–25). For nicotinamide (NAM, DOT Scientific, Burton, MI) treatment experiments, extracted parasites were first sorted from warm RPMI 1640 into room temperature RPMI 1640, and half of the parasites were placed in media supplemented with a final concentration of 250 μM NAM and incubated for 30 min. Heat-inactivated FBS (Gibco), 1 M NaCI (Thermo Fisher Scientific), and 1:1 3-methyl-1-butanol (Thermo Fisher Scientific) in mineral oil were used as cues. A curved platinum worm pick was used to remove L3s from warm media and place them on 0.8% agarose plates, plates were transferred to a 37°C incubator with 5% atmospheric CO_2_, and parasites were allowed to migrate for 30 minutes after which the plates were removed and scored. 8-10 parasites were added per plate. The chemotaxis index (CI) of each plate was calculated as follows: CI = (T - C) / (T + C + O), where T is the number of parasites that have migrated to the test cue, C is the number that migrate to the control cue, and O is the number that have migrated outside of the designated cue areas. To account for parasite injury in transfer, only plates that had C + T > 2 were used for statistical analysis and plotting. Experiments using FBS and NaCI as cues included 3 biological replicates, and experiments with 3-methyl-1-butanol included 2 biological replicates. Biological replicates are defined as groups of larvae originating from separate cohorts of mosquito infections (~250 infected mosquitoes) and separate cohorts of mf extractions. L3 stage parasites were subsequently extracted and assayed on different days. Technical variation was accounted for on each assay day by performing at least 3 assays (i.e. chemotaxis plates) per biological replicate.

### Larval coiling assay

NAM treatment of *B. pahangi* L3 was performed with parasites from in-house infections or received from the FR3. After extraction or receipt, parasites were washed with fresh RPMI and suspended in complete media (RPMI 1640 + 10% FBS + penicillin/streptomycin) at a density of 1 parasite per 2 μL. A 96-well plate was arranged with 50 μL of complete media with 2X NAM in each well, and parasites were pipetted into each individual well to create a density of 10-25 parasites per well in a final volume of 100 μL. Plates were immediately transferred to a 37°C incubator with 5% atmospheric CO_2_ and were left untouched until videos were recorded at 24 and 48 hours post-treatment. Care was taken not to disturb parasites while transferring the plates from the incubator to the recording stage. Parasites were allowed to cool at room temperature for 20 min. on the recording stage, after which each well was recorded for 10 sec. at 16 FPS. Recording was performed at 2.5X on a Zeiss Stemi 508 with a K LAB stand and oblique light with a monochrome CMOS camera (DMK 33UX178, The Imaging Source). Parasites were returned to the incubator after recording.

Coiling experiments included 3 biological replicates, defined as groups of larvae originating from separate cohorts of mosquito infections (~250 infected mosquitoes) and separate cohorts of mf extractions. L3 stage parasites were subsequently extracted and assayed on different days. An entire dose-response curve was performed for each replicate (100 nM to 1 mM), and at least 4 technical replicates (wells) of 10-25 parasites were included in each biological replicate.

Videos were manually scored and analyzed with a bespoke optical flow algorithm implemented in Python that calculates mean motility units (mmu) and has similarities to previous implementations (109,110). Relevant differences include the utilization of a dense flow algorithm that analyzes every pixel in the image instead of focusing on a sparse set of features; *post hoc* analysis rather than real time tracking to allow for greater quality control; image segmentation to calculate worm area to enable interwell normalization. Source code and a recommended Conda environment can be found at https://github.com/zamanianlab/BrugiaMotilityAnalysis.

For manual scoring of larval coiling, videos were assigned randomized file names and distributed to 3 researchers. Researchers blindly rated each well on a scale of 0-5 where 0 is the most coiling and 5 is the least coiling (the template with scoring instructions is provided in S3 File). Scores were collated and data was blindly analyzed and plotted with the tidyverse package (111) and custom R scripts.

### Aedes aegypti *feeding and engorgement assays*

3-4 day old adult female *Ae. aegypti* LVP were starved for 24 hours then provided with a blood meal for 30 minutes via a glass membrane feeder (n = 25-50) (83). Immediately after feeding, groups were cold anesthetized and visually inspected for distended abdomens to measure the proportion of feeding mosquitoes. Blood meal size was measured by calculating the ratio of the mosquito length (abdomen tip to thorax) to width (dorso-ventral width at the 5th abdominal segment) using Fiji (112)(S4 Figure).

### Larval temperature-shift assay and qPCR

*B. pahangi* L3s were extracted in bulk and separated into three treatment treatment groups: immediate storage in Trizol LS (Ambion); 1 mL RPMI 1640 + 10% FBS + penicillin/streptomycin at room temperature; or 1 mL RPMI 1640 + 10% FBS + penicillin/streptomycin in a 37°C heat block. Parasites were incubated in media for 4 hours. After incubation, media was removed, parasites were washed once in fresh RPMI 1640 and stored in Trizol LS at −80°C until processing. To extract RNA, samples were thawed on ice, and the volume was adjusted to a final ratio of 3 Trizol LS: 1 RNase-free water. Samples were lysed with a TissueLyser LT (Qiagen, Venlo, The Netherlands). One 5 mm stainless steel bead was added to each tube, which then underwent two cycles of 3 minutes of shaking at 30 Hz. Tubes were cooled on ice for 2 minutes in between cycles. RNA was extracted with the Direct-zol RNA miniprep kit (Zymo) according to the manufacturer’s instructions, including an on-column DNase treatment, and RNA was eluted in 15 μL RNase-free water. RNA samples were quantified with a NanoDrop 1000 and immediately used for first-strand cDNA synthesis with SuperScript III (Thermo Fisher Scientific) using random hexamers and normalizing RNA input. cDNA was stored at −20°C until further use.

For qPCR, GAPDH control primers(113) and *osm-9* primers (designed with Primer3(114), F: CCCGCTGATCCAAACATTG, R: TGCACTACACGTCATATCACTG) were optimized with *B. pahangi* L3 RNA from the FR3 with cDNA synthesized using the same SuperScript III master mix as the experimental RNA samples. A 20 μL reaction was used with 2X PowerUp SYBR Green MasterMix, 800 nM primers, and 5.2 ng RNA. Reactions were run in duplicate on a StepOnePlus real-time PCR system. C_T_ values were calculated with the system’s automatic threshold, and relative expression was calculated with the ΔΔC_T_ method(87).

### In squito *exposure of infective larva to Bpa-osm-9 and Bpa-tax-4 dsRNAs*

Primers were designed to amplify 200-600 bp regions from cloned *Bma-tax-4* and *Bma-osm-9* which had >95% identity with their *B. pahangi* orthologs, and T7 recognition sequences were appended to the 5’ end of each primer. Cloned genes (below) were used as template DNA for PCRs with Phusion polymerase (New England Biolabs, Ipswich, MA). Complete dsRNA synthesis protocols, including primer sequences and thermocycler programs, can be found in S3 File. PCR product was cleaned (Qiagen MinElute PCR Purification Kit) and resuspended in water at a desired concentration of 1-2 ug/uL as measured by a Qubit 3.0 dsDNA assay (Thermo Fisher Scientific). This product was subsequently used as the template for a dsRNA synthesis reaction (MegaScript RNAi, Thermo Fisher Scientific). dsRNA was DNase treated, purified with phenol/chloroform, and resuspended in nuclease-free water at a concentration of 1-4 ug/uL. The concentrations and purity of 1:20 dilutions of dsRNA were measured with a NanoDrop 1000 (Thermo Fisher Scientific).

*Ae. aegypti* LVP were infected in batches of 250 with *B. pahangi* mf as described above. After blood feeding, 25 mosquitoes were organized into small cardboard cartons in preparation for injection. Injections of dsRNA were carried out with the following modifications of an established protocol (63). Prior to injection, infected mosquitoes were starved by removing sucrose pads 8 DPI. At 9 DPI, infected mosquitoes were injected with 250 μL of 1 ug/uL dsRNA, coinciding with the L2 to L3 molt in the thoracic musculature(8). Mosquitoes were injected in cohorts of 25, and cohorts were immediately returned to 26°C and sucrose pads were replaced. Dead mosquitoes were removed daily until time of assay at 14 DPI, at which point mosquitoes were dissected to extract L3s for use in chemotaxis assays.

### *Long-read sequencing in* Brugia malayi *adult males and females*

Total RNA from *B. malayi* adult males and females was obtained from the FR3. RNA quality was assessed by 2100 Bioanalyzer, converted to single stranded cDNA and amplified using the SMARTer PCR cDNA Synthesis Kit (Takara Bio, Kusatsu, Japan), and IsoSeq libraries were constructed with equimolar cDNA fractions (0.5X and 1X) with the SMRTbell Template Prep Kit 1.0 (Pacific Biosciences, Menlo Park, CA). Library quantity and quality were assessed by Qubit HS DNA (Thermo Fisher Scientific) and 2100 Bioanalyzer. Isoforms were clustered and polished from subreads with IsoSeq2, visualized with IGV(115), and annotated with BLAST(116).

### Cloning of osm-9 and tax-4 homologs

Primers directed toward the ATG start, stop codon, or 3’ UTR region of the IsoSeq-generated gene model of *Bma-osm-9* and the predicted gene models of *Bma-tax-4* and *Bma-ocr-1/2a* (the *B. malayi* gene with the highest amino acid identity to *Cel-ocr-2*) were designed with Primer3(114). A full length amplicon was produced with Phusion or Q5 polymerases (New England Biolabs). Amplicons were A-tailed with GoTaq Flexi (Promega, Madison, WI) and cloned into pGEM-T in JM109 competent cells (Promega). *C. elegans* N2 genomic DNA was extracted with the Qiagen DNeasy kit. A ~1.6 kb portion upstream of *Cel-osm-9(30)a* ~3 kb portion upstream of *Cel-tax-4(28*) were amplified and cloned into pGEM-T as above. Final expression constructs were assembled with the HiFi Assembly kit (New England Biolabs), using amplicons generated with Q5 polymerase from the promoter and gene as two fragments, and pPD95.75 (a gift from Andrew Fire (Addgene plasmid # 1494; http://n2t.net/addgene:1494; RRID:Addgene_1494)) double-digested with Xbal and EcoRI, as the backbone, and a BamHI restriction site was added between the promoter and genes. *C. elegans osm-9* and *tax-4* were amplified from plasmids (gifts from Shawn Xu (117) and Cornelia Bargmann (118), respectively) with Q5 polymerase. Each *C. elegans* gene was assembled into previously created expression vectors by replacing *B. malayi* genes with BamHI/EcoRI double digestions. The *unc-54* 3’ UTR of the pPD95.75 backbone was replaced in all constructs containing *osm-9* homologs by double-digesting final constructs with EcoRI/BsiWI or PCR amplifying the entire plasmid without the *unc-54* 3’ UTR and assembling the resulting fragments with the *Cel-osm-9* 3’ UTR amplicon. Complete cloning protocols, including primer sequences and thermocycler programs, can be found in S4 File. All products were verified with Sanger sequencing. Expression vectors were injected into *C. elegans* hermaphrodites as described and transgenic strains were used for rescue experiments.

### Caenorhabditis elegans *sensory assays*

Population chemotaxis assays were performed as described(119). For each strain, five L4 worms from three independently derived lines were picked to each of five seeded NGM plates 5 days before the assay date. On assay day, worms from independent lines were washed off plates and pooled with M9 into a single tube per strain, washed with M9 three times and once with water. For each of five 10 cm chemotaxis plates (2% agar, 5 mM KH_2_PO_4_/K_2_HPO_4_ pH 6.0, 1 mM CaCl_2_ and 1 mM MgSO_4_) per strain, 1 μL of 1 M sodium azide was place on opposite sides of the plate and allowed to soak in with the plate lids removed. Once dry, 1 μL of cue and diluent were then placed at the same location as the sodium azide on opposing sides of the plate. 1:1000 diacetyl (Santa Cruz Biotechnology, Santa Cruz, CA) and 1:10 isoamyl alcohol (Thermo Fisher Scientific) in ethanol were used as cues for the *osm-9* and *tax-4* experiments, respectively. After the addition of cues, 100-200 worms were quickly pipetted to the center of assay plates. Excess water was removed with a Kimwipe (Kimberly-Clark, Irving, TX), and worms were gently spread with a platinum worm pick. Plates were left untouched on a benchtop for 60 minutes at room temperature (~21°C), after which animals were counted in total and at each cue and control region. The chemotaxis index (CI) of each plate was calculated as follows: CI = (T - C) / (T + C + O), where T is the number of worms that were paralyzed at the test cue, C is the number at the control, and O is the number that had migrated to neither the test nor the cue.

Benzaldehyde avoidance assays were performed as described (30,120). Young adult animals were transferred to unseeded NGM plates and 2 μL of benzaldehyde (Sigma-Aldrich) in a 20 μL borosilicate capillary was held in front of the animal’s nose and the time to reversal was recorded. Five animals per strain were exposed a single time for each replicate and a minimum of 3 replicates were performed.

Nose-touch reversal assays were performed as described(121). Young adult animals were transferred to unseeded NGM plates and observed for reversal movement after colliding head-on with an eyelash. For each replicate, five animals were strain were observed for reversal after 10 successive collisions, and a minimum of 3 replicates were performed.

## Acknowledgements

Some *C. elegans* strains were provided by the CGC, which is funded by the NIH Office of Research Infrastructure Programs (P40 OD010440). Some parasite materials were provided by the NIH/NIAID Filariasis Research Reagent Resource Center (www.filariasiscenter.org). Sanger sequencing and RNA-seq was carried out at the University of Wisconsin-Madison Biotechnology Center. Additionally, the authors would like to thank Tran To and Elena Garncarz for their assistance with the *C. elegans* behavioral assays.

## Supporting Information

**S1 Table.** Spreadsheet of species included in the comparative analysis, clade designation, genome BioProject, and whether or not each species is represented in the trees

**S1 Data.** Chemoreceptor IQ-TREE consensus tree in Newick format

**S2 Data.** TRP MrBayes consensus tree in Nexus format

**S3 Data.** CNG MrBayes consensus tree in Nexus format

**S4 Data.** List of *C. elegans* chemoreceptors IDs

**S1 Figure.** Flow chart of comparative genomics pipeline

**S2 Figure.** Total chemoreceptor count as a function of genome contiguity.

**S3 Figure.** Alternative plot of *B. malayi* head/tail RNA-seq

**S4 Figure.** Motility analysis of L3 parasites after cooling and warming.

**S5 Figure.** Representative images of measured mosquito abdomens

**S6 Figure.** L3 recovery correlation plot

**S7 Figure.** osm-9 nucleotide alignment

**S8 Figure.** osm-9 amino acid alignment, including Inactive from *Drosophila melanogaster*

**S9 Figure.** tax-4 amino acid alignment

**S10 Figure.** qPCR results for knock-out and transgenic strains. ND = “not determined.”

**S1 File.** List of all identified chemoreceptors with family and superfamily annotations

**S2 File.** List of nematode species included in Figure 1C with assigned category and justification

**S3 File.** Template with instructions for assigning L3 coiling scores.

**S4 File.** Complete protocols for all cloning efforts

