## Supplementary material for "Genetic and functional diversification of chemosensory pathway receptors in mosquito-borne filarial nematodes": S1 Figure

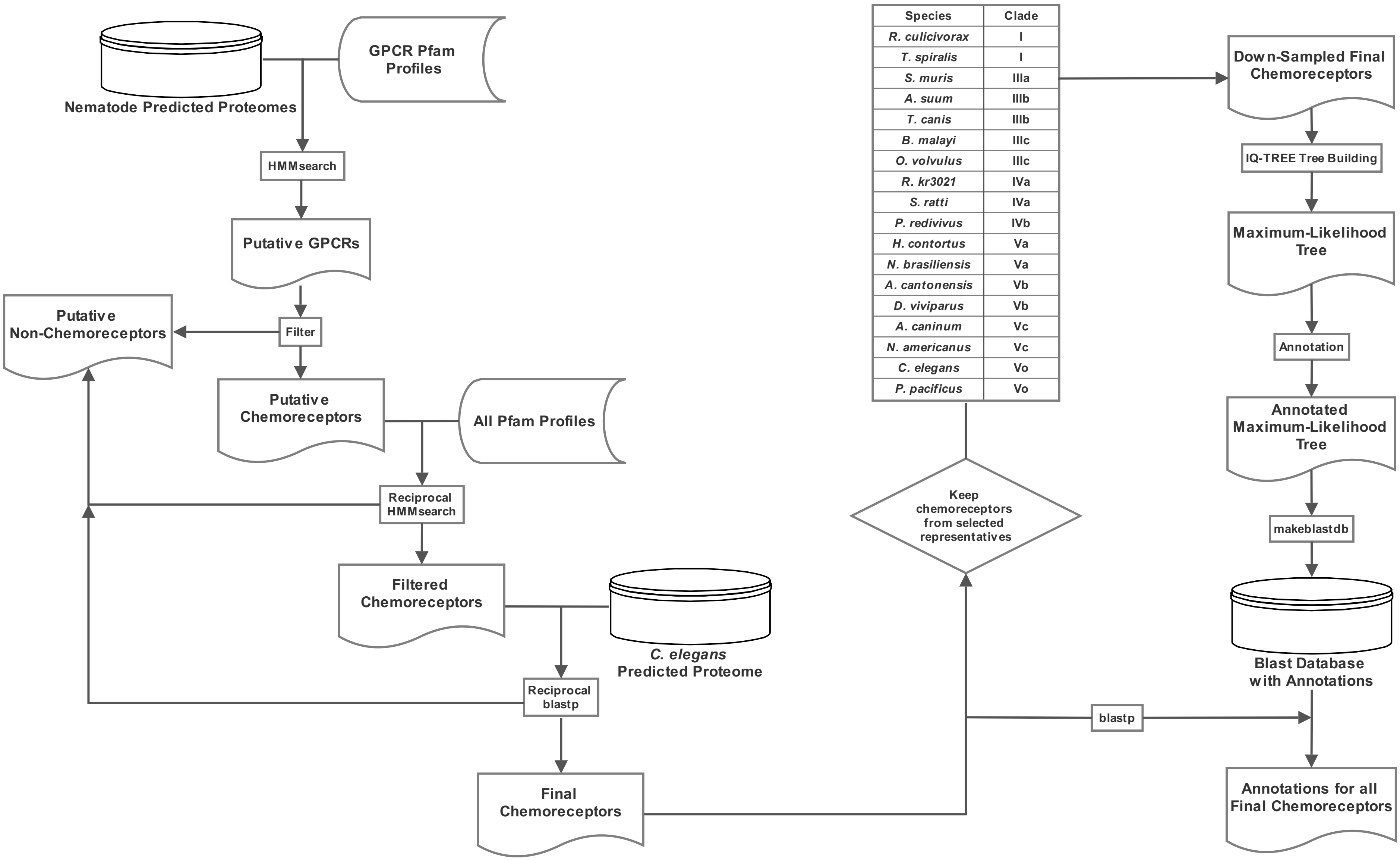


| Species | Clade |
|---|---|
| *R. culicivorax* | I |
| *T. spiralis* | I |
| *S. muris* | IIIa |
| *A. suum* | IIIb |
| *T. canis* | IIIb |
| *B. malayi* | IIIc |
| *O. volvulus* | IIIc |
| *R. kr3021* | IVa |
| *S. ratti* | IVa |
| *P. redivivus* | IVb |
| *H. contortus* | Va |
| *N. brasiliensis* | Va |
| *A. cantonensis* | Vb |
| *D. viviparus* | Vb |
| *A. caninum* | Vc |
| *N. americanus* | Vc |
| *C. elegans* | Vo |
| *P. pacificus* | Vo |
