## Supplementary figures and images for "Genetic and functional diversification of chemosensory pathway receptors in mosquito-borne filarial nematodes"

### S2 Figure

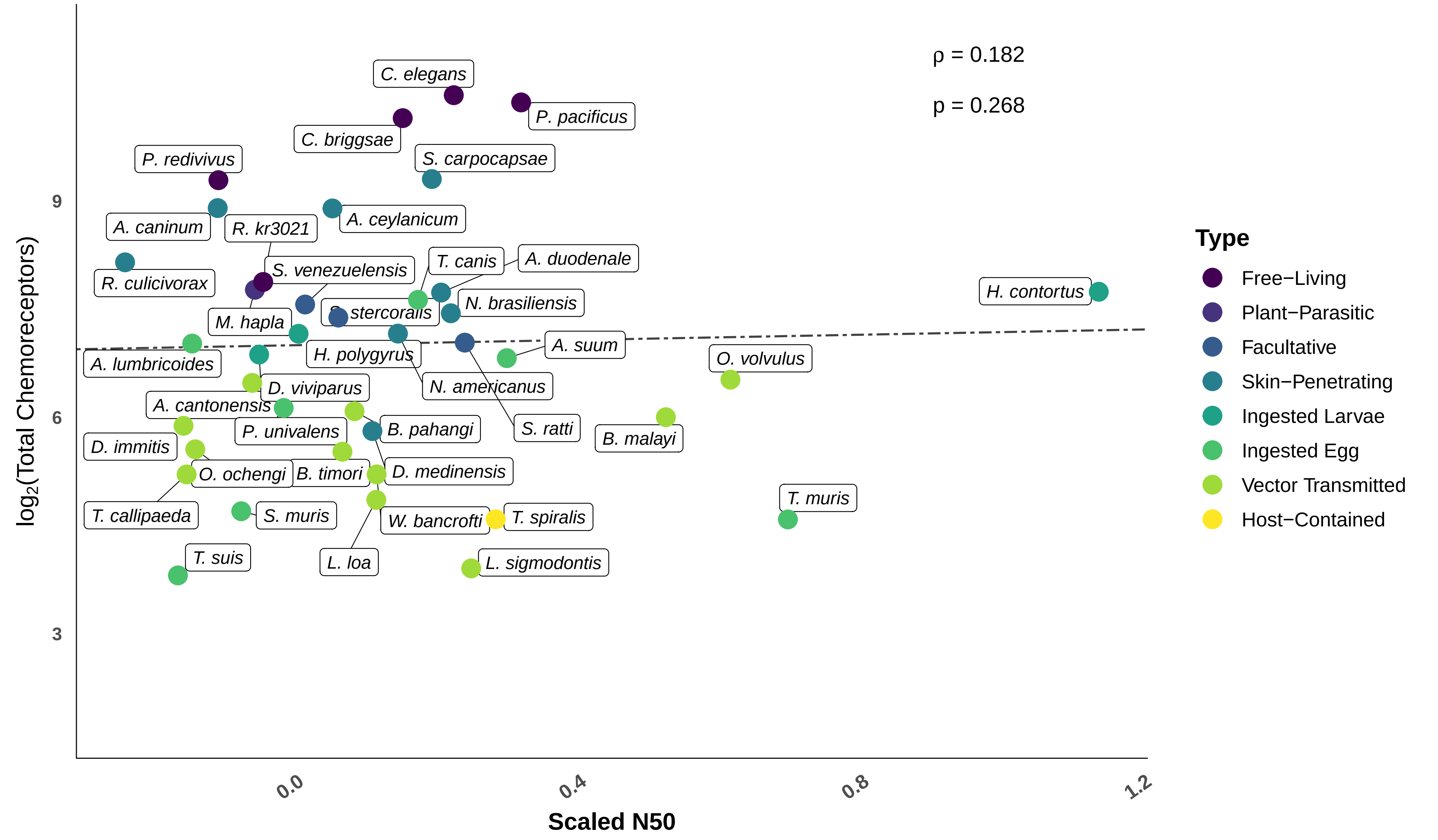

### S3 Fig

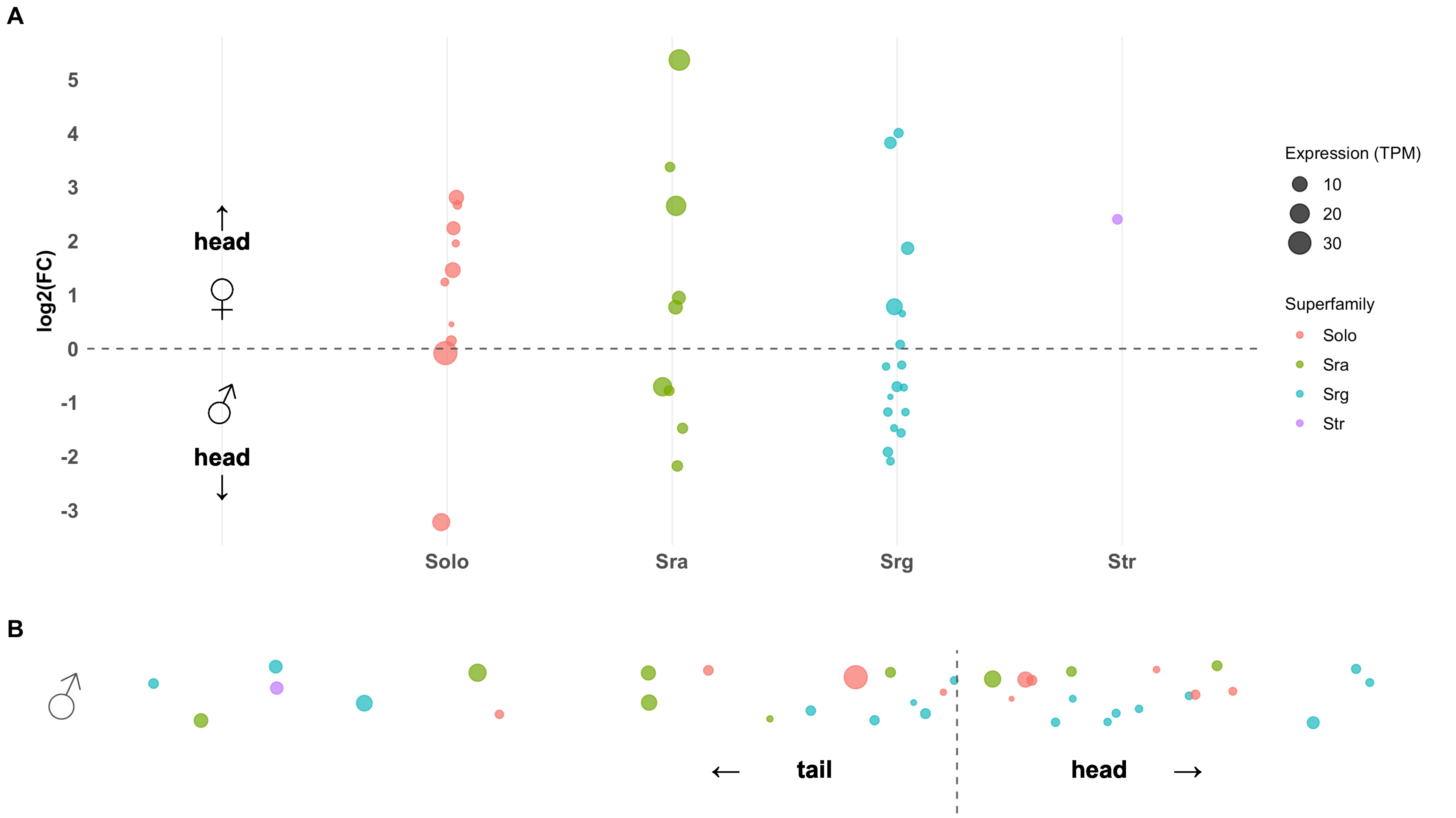

### S4 Figure

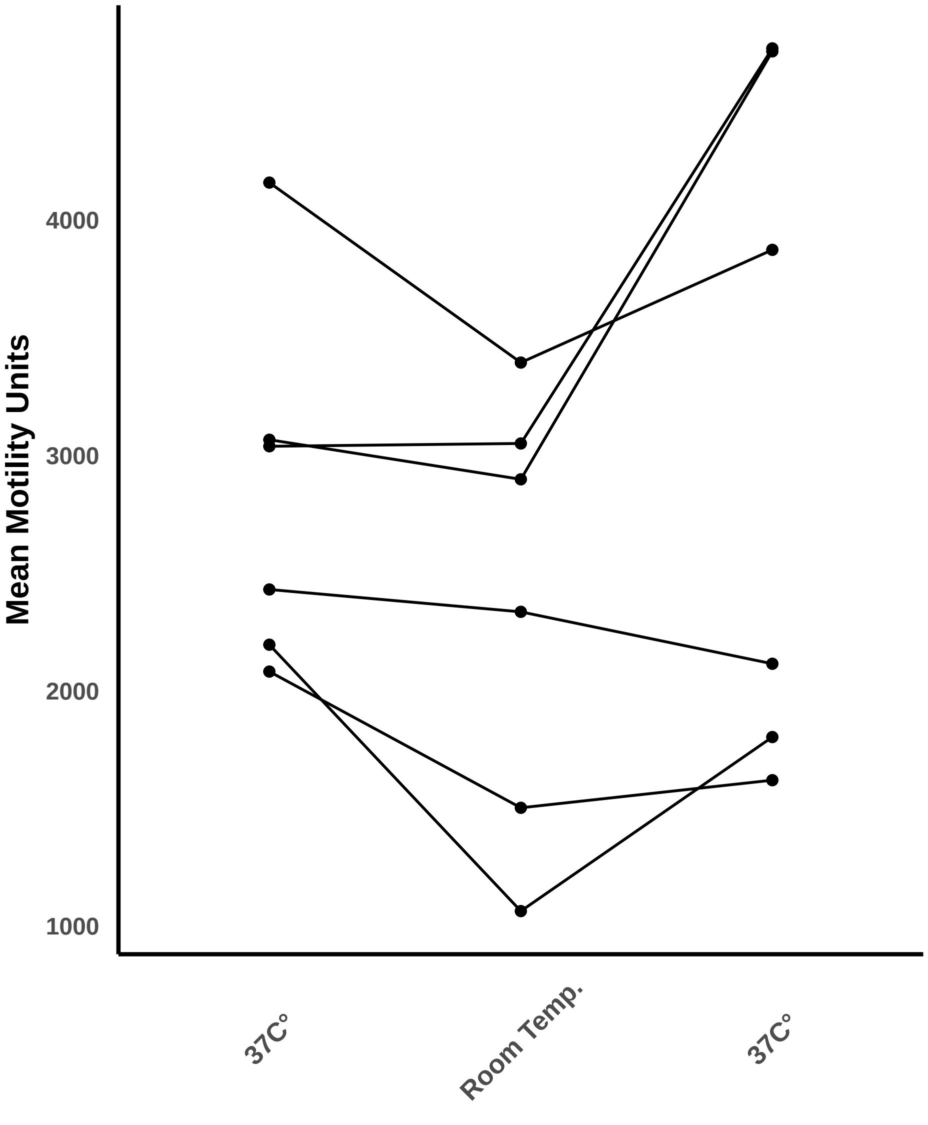

### S5 Figure

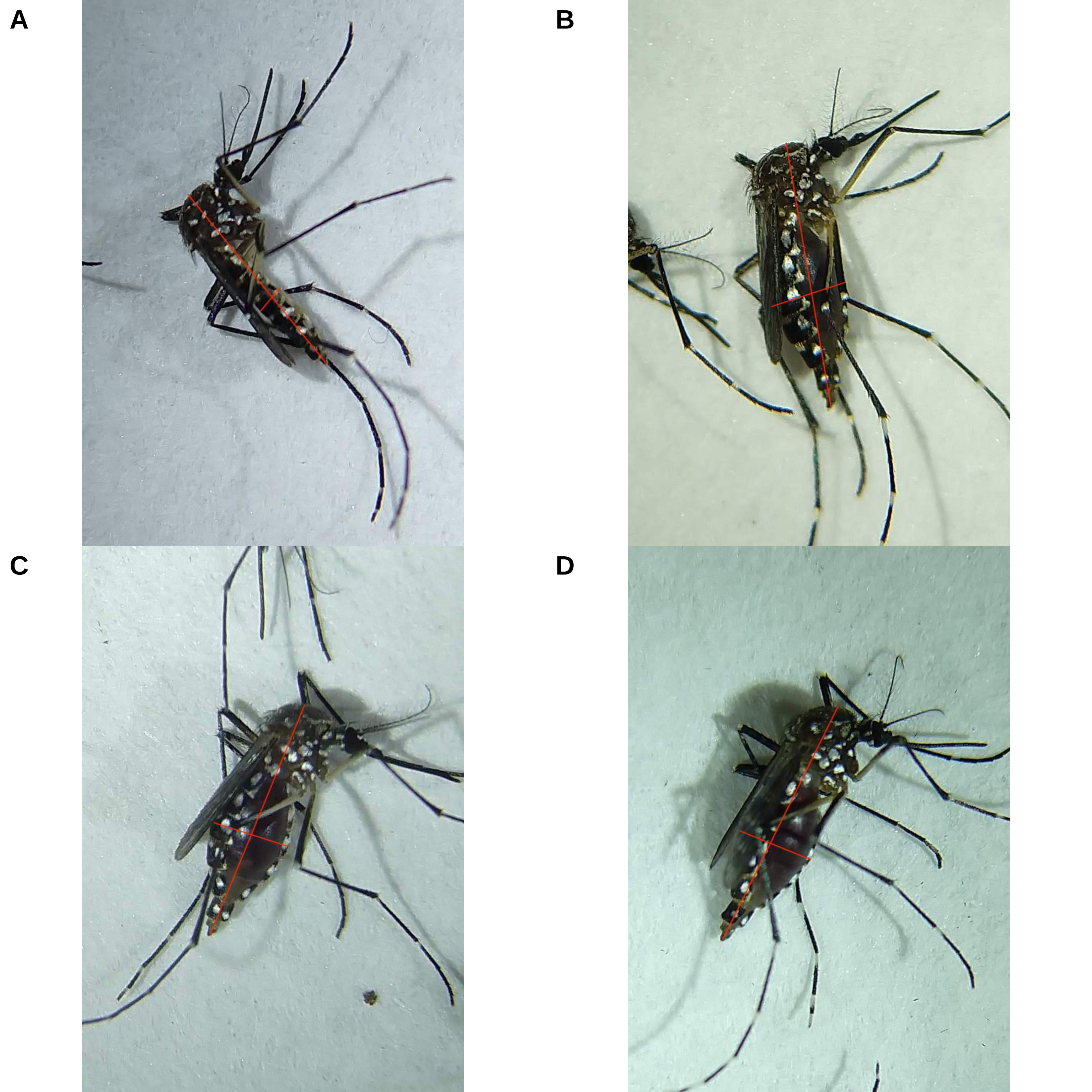

### S6 Figure

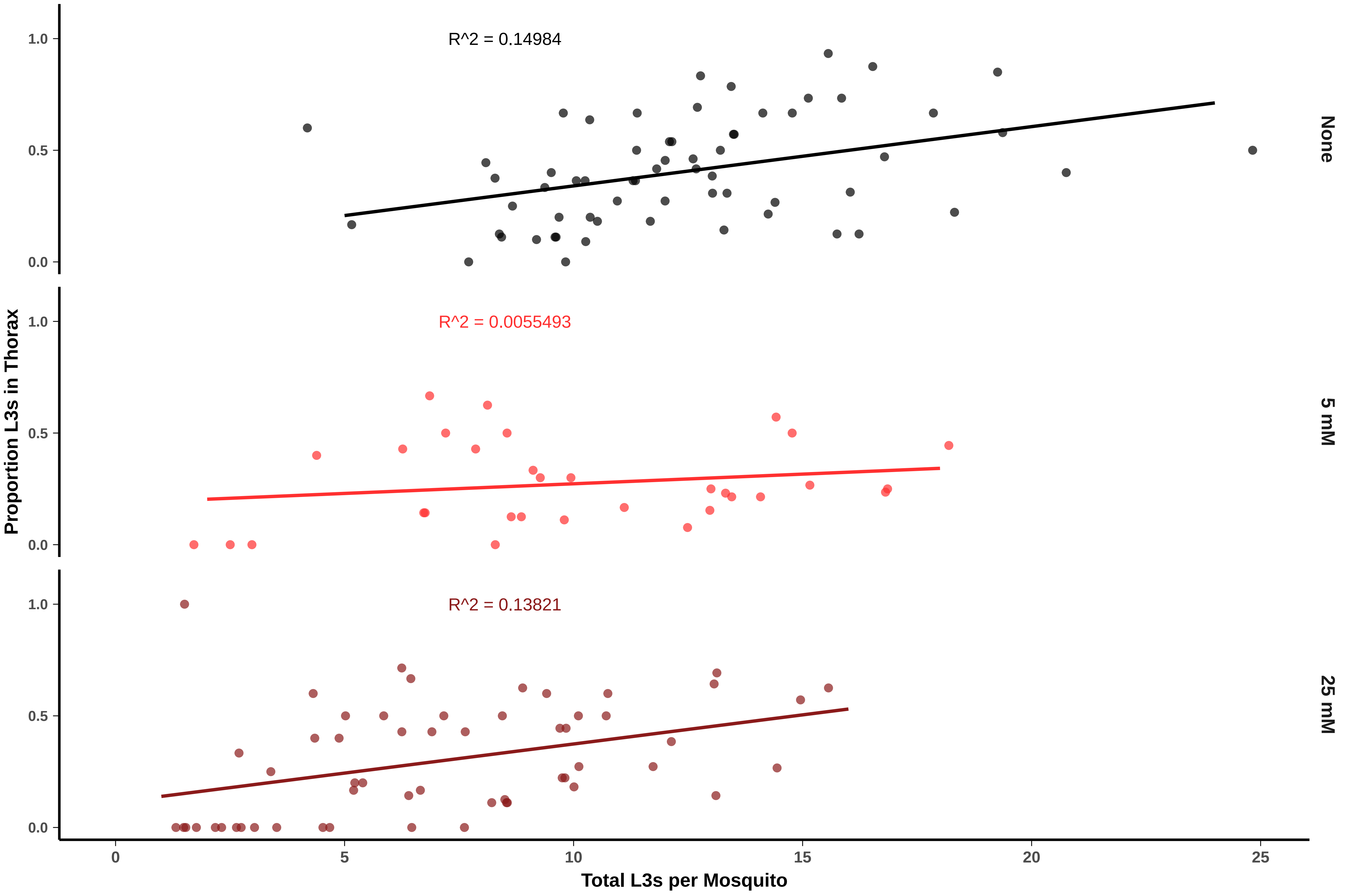
