## Supplementary material for "Genetic and functional diversification of chemosensory pathway receptors in mosquito-borne filarial nematodes": S4 File

### Cloning, dsRNA Synthesis, and qPCR Protocols

#### *Brugia malayi* Gene Cloning

##### *Bm-osm-9* expression cloning

**ORF cloning:**

Forward primer: ATGGGACAACTAAAGAGCAAAA
Reverse primer: TCAGCCACTGAAATTGAA

Template: Adult female *Brugia malayi* cDNA synthesized with oligo(dT) primers

| Stage | Temperature | Length | Cycles |
|---|---|---|---|
| Melt | 98°C | 30 s | |
| Melt | 98°C | 10 s | |
| Anneal | 58°C | 30 s | 40 |
| Extend | 65°C | 3 min | |
| Extend | 65°C | 5 min | |

Primers for Sanger sequencing:

M13F
M13R
TACAATATGGCGCTGATCCA
CGTATATCGGGCAATGTTCA

**Promoter cloning:**

Forward primer: GGTGGT-TCTAGA-TTTTAAAAAGGTTTTTGAGAATCAG
Reverse primer: GGTGGT-GGATCC-GTTTGGTTTCTGAAAAAATTGG

Template: *C. elegans* N2 genomic DNA

| Stage | Temperature | Length | Cycles |
|---|---|---|---|
| Melt | 98°C | 30 s | |
| Melt | 98°C | 10 s | |
| Anneal | 63°C | 30 s | 5 |
| Extend | 65°C | 1:40 min | |
| Melt | 98°C | 10 s | |
| Anneal/Extend | 65°C | 2:10 min | 30 |
| Extend | 65°C | 5 min | |

##### Preparation for HiFi Assembly

##### *Ce-osm-9* promoter

Forward primer: TGCATGCCTGCAGGTCGACTCTAGA-TTTTAAAAAGGTTTTTGAGAATCAG
Reverse primer: GTTGTCCCAT-GGATCC-GTTTGGTTTCTGAAAAAATTG

Template: *Ce-osm-9* promoter in pGEM-T

| Stage | Temperature | Length | Cycles |
|---|---|---|---|
| Melt | 98°C | 30 s | |
| Melt | 98°C | 10 s | |
| Anneal | 59°C | 30 s | 35 |
| Extend | 72°C | 1:30 min | |
| Extend | 65°C | 5 min | |

##### *Bm-osm-9* ORF

Forward primer: GAAACCAAACGGATCC-ATGGGACAACTAAAGAGC
Reverse primer: AGCGACCGGCGCTCAGTTG-GAATTC-AGCCACTGAAATTGAAG

Template: *Bm-osm-9* ORF in pGEM-T

| Stage | Temperature | Length | Cycles |
|---|---|---|---|
| Melt | 98°C | 30 s | |
| Melt | 98°C | 10 s | |
| Anneal | 67°C | 30 s | 35 |
| Extend | 72°C | 1:30 min | |
| Extend | 65°C | 5 min | |

##### HiFi Assembly Parameters

Vector amount: 0.023 pmol
Promoter amount: 0.046 pmol
ORF amount: 0.046 pmol

Primers for Sanger sequencing:

CGATGGATACGCTAACAACTTGG
ACTAAGAAGGCGGAGTTGGC
TACAATATGGCGCTGATCCA
CGTATATCGGGCAATGTTCA
GAGCACAGGGAGAAAGAGCA

##### *Bm-ocr-1/2a* expression cloning

##### ORF cloning:

Forward primer: ATGGGAAATGTGGAATCGTCG
Reverse primer: TTGTCGATGCCACAGTGAAC

Template: Adult female *Brugia malayi* cDNA synthesized with random hexamer primers

| Stage | Temperature | Length | Cycles |
|---|---|---|---|
| Melt | 98°C | 30 s | |
| Melt | 98°C | 10 s | |
| Anneal | 58°C | 30 s | 40 |
| Extend | 65°C | 3 min | |
| Extend | 65°C | 5 min | |

Primers for Sanger sequencing:

M13F
M13R
TCTACTTCGACATGGTGCTGA
GAGCACATCAAACGAAATGGT

**Preparation for HiFi Assembly**

***Bm-ocr-1/2a* ORF**

Forward primer: ATTTTTTCAGAAACCAAAC-GGATCC-ATGGGAAATGTGGAATCG
Reverse primer: AGCGACCGGCGCTCAGTTG-GAATTC-TATTCTCCTAGAACCTTTAATC

Template: *Bm-ocr-1/2a* ORF in pGEM-T

| Stage | Temperature | Length | Cycles |
|---|---|---|---|
| Melt | 98°C | 30 s | |
| Melt | 98°C | 10 s | |
| Anneal | 58°C | 30 s | 35 |
| Extend | 72°C | 1:30 min | |
| Extend | 72°C | 5 min | |

**HiFi Assembly Parameters**

Vector amount: 0.023 pmol
Promoter amount: 0.046 pmol
ORF amount: 0.046 pmol

Primers for Sanger sequencing:

ACTAAGAAGGCGGAGTTGGC
TCTACTTCGACATGGTGCTGA
GAGCACATCAAACGAAATGGT
GAGCACAGGGAGAAAGAGCA

***Bm-tax-4* expression cloning**

**ORF cloning:**

Forward primer: ATGTTCTCTAAAAGTCATGATGA
Reverse primer: TTAACTATCACATATCATCTGATAATC

Template: Adult female *Brugia malayi* cDNA synthesized with oligo(dT) primers

| Stage | Temperature | Length | Cycles |
|---|---|---|---|
| Melt | 98°C | 30 s | |
| Melt | 98°C | 10 s | |
| Anneal | 58°C | 30 s | 40 |
| Extend | 65°C | 1:30 min | |
| Extend | 65°C | 5 min | |

Primers for Sanger sequencing:

M13F
M13R
CCTGAAAAATGCGTACTCTTCTAA

**Promoter cloning:**

Forward primer: GGTGGT-TCTAGA-ACCATCACTGAAGGGTGAGC
Reverse primer: GGTGGT-GGATCC-TCTTGA-AACATAATTAAATTTTGAGAATGATAG

Template: *C. elegans* N2 genomic DNA

| Stage | Temperature | Length | Cycles |
|---|---|---|---|
| Melt | 98°C | 30 s | |
| Melt | 98°C | 10 s | |
| Anneal | 64°C | 30 s | 5 |
| Extend | 65°C | 1:40 min | |
| Melt | 98°C | 10 s | |
| Anneal/Extend | 65°C | 2:10 min | 30 |
| Extend | 65°C | 5 min | |

**Preparation for HiFi Assembly**

***Ce-tax-4* promoter**

Forward primer: TGCATGCCTGCAGGTCGACTCTAGAGGTGGTTCTAGA-ACCATCACTG
Reverse primer: TAGAGAACA-TGGTGGT-GGATCC-TCTTGAAAC

Template: *Ce-tax-4* promoter in pGEM-T

| Stage | Temperature | Length | Cycles |
|---|---|---|---|
| Melt | 98°C | 30 s | |
| Melt | 98°C | 10 s | |
| Anneal | 65°C | 30 s | 35 |
| Extend | 72°C | 1:30 min | |
| Extend | 72°C | 5 min | |

##### *Bm-tax-4* ORF

Forward primer: ATCCACCACC-ATGTTCTCTAAAAGTCATGATGATAC
Reverse primer: AGCGACCGGCGCTCAGTTGGAATTC-TTAACTATCACATATCATCTGATAATC

Template: *Bm-tax-4* ORF in pGEM-T

| Stage | Temperature | Length | Cycles |
|---|---|---|---|
| Melt | 98°C | 30 s | |
| Melt | 98°C | 10 s | |
| Anneal | 59°C | 30 s | 35 |
| Extend | 72°C | 1:30 min | |
| Extend | 72°C | 5 min | |

##### HiFi Assembly Parameters

Vector amount: 0.023 pmol
Promoter amount: 0.046 pmol
ORF amount: 0.046 pmol

Primers for Sanger sequencing:

CGATGGATACGCTAACAACTTGG
TAGAGAACATGGTGGTGGATCCTCTTGAAAC
CGTAAATAGGGTATTGATCGTTGA
CCTGAAAAAATGCGTACTCTTCTAA
GAGCACAGGGAGAAAGAGCA

#### *Caenorhabditis elegans* Gene Cloning

##### *Ce-osm-9* expression cloning

##### Preparation for HiFi Assembly

Forward primer: ATTTTTTCAGAAACCAAAC-GGATCC-ATGGGCGGTGGAAGTTCG
Reverse primer: AGCGACCGGCGCTCAGTTG-GAATTC-TCATTCGCTTTTGTCATTTGTCG

Template: Psra-6::osm-9(cDNA)::sl2::CFP vector from the Shawn Xu lab (Neuron, Wang 2016)

| Stage | Temperature | Length | Cycles |
|---|---|---|---|
| Melt | 98°C | 30 s | |
| Melt | 98°C | 10 s | |
| Anneal | 65°C | 30 s | 35 |
| Extend | 72°C | 1:10 min | |
| Extend | 72°C | 2 min | |

**HiFi Assembly Parameters**

Vector amount: 0.015 pmol
ORF amount: 0.046 pmol

Primers for Sanger sequencing:

ACTAAGAAGGCGGAGTTGGC
GCATTTGCCGCTTGTTTTGG
CGGCGCCAACAACCATTG
GAGCACAGGGAGAAAGAGCA

**_Ce-tax-4_ expression cloning**

**Preparation for HiFi Assembly**

Forward primer: ATTTAATTATGTTTCAAGAG-GGATCC-ATGTCAACGGCGGAACCTG
Reverse primer: ACCGGCGCTCAGTTGGAATT-GAATTC-CTATTTGAGCAAGGATTCAGATTCAGTTC

Template: pEM04 (gcy-36p::tax-4::sl2::GFP) vector from the Cornelia Bargmann lab (Nature, Macosko 2009)

| Stage | Temperature | Length | Cycles |
|---|---|---|---|
| Melt | 98°C | 30 s | |
| Melt | 98°C | 10 s | |
| Anneal | 66°C | 30 s | 35 |
| Extend | 72°C | 1:15 min | |
| Extend | 72°C | 2 min | |

**HiFi Assembly Parameters**

Vector amount: 0.012 pmol
ORF amount: 0.024 pmol

Primers for Sanger sequencing:

TTTGTGGTTCAGGCTGCTCA

TCAACGATCTCATTGGACCATCT
TCCGGTGCATTTTCATCCAA
GAGCACAGGGAGAAAGAGCA

#### Replacing *unc-54* 3' UTR with *osm-9* 3' UTR

##### ***Bm-osm-9*** **expression cloning**

##### Preparation for HiFi Assembly

###### ***osm-9*** **3' UTR cloning:**

Forward primer: AACTTCAATTTCAGTGGCTGAATTC-GAACTTTTTTCTTCTAATTTTTTGAAAAC
Reverse primer: CGCGCGAGACGAAAGGGCCCGTACG-AGTAAATTTGGCAATTTCTG

Template: *C. elegans* N2 genomic DNA

| Stage | Temperature | Length | Cycles |
|---|---|---|---|
| Melt | 98°C | 30 s | |
| Melt | 98°C | 10 s | |
| Anneal | 56°C | 30 s | 35 |
| Extend | 72°C | 2 min | |
| Extend | 72°C | 2 min | |

###### HiFi Assembly Parameters

Vector amount: 0.0115 pmol
ORF amount: 0.0231 pmol

Primers for Sanger sequencing:

GTTTCGGCGCAAAGACAGTT
TTAAAGGGCGCACTCTTCCG
TCCCGAAAGTTGATCTCCGA
AACTTTTGGAGGCGGGTGAG
CATAGTTAAGCCAGCCCCGA

##### ***Ce-osm-9*** **expression cloning**

##### Preparation for HiFi Assembly

###### ***osm-9*** **3' UTR cloning:**

Forward primer: ACAAAAGCGAATGAGAATTCGAACTTTTTTCTTC-TAATTTTTTGAAAACT
Reverse primer: GAGACGAAAGGGCCCGTACGAGTAAATTTG-GCAATTTCTGGC

Template: *C. elegans* N2 genomic DNA

| Stage | Temperature | Length | Cycles |
|---|---|---|---|
| Melt | 98°C | 30 s | |
| Melt | 98°C | 10 s | |
| Anneal | 60°C | 30 s | 35 |
| Extend | 72°C | 2 min | |
| Extend | 72°C | 2 min | |

***Ce-osm-9* backbone cloning:**

Forward primer: CAGAAATTGCCAAATTTACTCGTACGGGCC-CTTTCGTCTC
Reverse primer: AAATTAGAAGAAAAAAGTTCGAATTCTCATTCGCTTTTGT-CATTTG

Template: mazEx18

| Stage | Temperature | Length | Cycles |
|---|---|---|---|
| Melt | 98°C | 30 s | |
| Melt | 98°C | 10 s | |
| Anneal | 64°C | 30 s | 35 |
| Extend | 72°C | 3:30 min | |
| Extend | 72°C | 2 min | |

**HiFi Assembly Parameters**

Vector amount: 0.0114 pmol
ORF amount: 0.0228 pmol

Primers for Sanger sequencing:

CCGGCTTCGTTTGATCAGAG
TTAAAGGGCGCACTCTTCCG
TCCCGAAAGTTGATCTCCGA
AACTTTTGGAGGCGGGTGAG
CATAGTTAAGCCAGCCCCGA

#### dsRNA Synthesis

##### *Bm-osm-9* Target

Forward primer + T7: CGATGTTAATACGACTCACTATAGGG-CACCATTGACGCTTGCAACA
Reverse primer + T7: CGATGTTAATACGACTCACTATAGGG-ACCGCACCCAATCATTCCAT

Template: *Bm-osm-9* Expression Plasmid

| Stage | Temperature | Length | Cycles |
|---|---|---|---|
| Melt | 98°C | 30 s | |
| Melt | 98°C | 10 s | |
| Anneal | 59°C | 30 s | 35 |
| Extend | 72°C | 1:30 min | |
| Extend | 65°C | 5 min | |

##### ***Bm-tax-4*** **Target**

Forward primer + T7: TAATACGACTCACTATAGGGAGA-TTCCGGATAAATTGCAAACAG
Reverse primer + T7: TAATACGACTCACTATAGGGAGA-CAAATCGAGCCATTTTGGTT

Template: *Bm-tax-4* Expression Plasmid

| Stage | Temperature | Length | Cycles |
|---|---|---|---|
| Melt | 98°C | 30 s | |
| Melt | 98°C | 10 s | |
| Anneal | 59°C | 30 s | 35 |
| Extend | 72°C | 1:30 min | |
| Extend | 65°C | 5 min | |

#### qPCR Primers

| Target | Sequence |
|---|---|
| Bm-GAPDH | TTTCTGCAGAGGGAGGCAAG |
| Bm-GAPDH | TCAGCGGGATCTTTGCTGTT |
| Bm-osm-9 | CCCGCTGATCCAAACATTG |
| Bm-osm-9 | TGCACTACACGTCATATCACTG |
| Bm-tax-4 | TTGGCTCAGATGGTTGGGTC |
| Bm-tax-4 | TTGGACTTGGCACTTCACCG |
| Bm-ocr-1/2a | ATCTACGTTGGCAAGGCGAT |
| Bm-ocr-1/2a | CCACTGTCGTTTCCACTCAG |
| Y45F10D.4 | GTCGCTTCAAATCAGTTCAGC |
| Y45F10D.4 | GTTCTTGTCAAGTGATCCGACA |
