## Supplementary material for "Genetic and functional diversification of chemosensory pathway receptors in mosquito-borne filarial nematodes": S7 Figure

```
                                  1
Bm-osm-9_Bm1711.1                 ATGGGACAAC TAAAGAGCAA AATTTTGCAC CAAACTGGAG GAGAAGATAT TGACAATTTA AAAGAAGATC CTGACGATCA ATGGTCGAAT TTATATCGTG AACGTAAAAA GAATCACCTT
Bm-osm-9_clone                    ATGGGACAAC TAAAGAGCAA AATTTTGCAC CAAACTGGAG GAGAAGATAT TGACAATTTA AAAGAAGATC CTGACGATCA ATGGTCGAAT TTATATCGTG AACGTAAAAA GAATCACCTT
i3_LQ_NR42497|c19820/f1p1/3020    ATGGGACAAC TAAAGAGCAA AATTTTGCAC CAAACTGGAG GAGAAGATAT TGACAATTTA AAAGAAGATC CTGACGATCA ATGGTCGAAT TTATATCGTG AACGTAAAAA GAATCACCTT
i2_LQ_NR42497|c51951/f1p6/3009    ATGGGACAAC TAAAGAGCAA AATTTTGCAC CAAACTGGAG GAGAAGATAT TGACAATTTA AAAGAAGATC CTGACGATCA ATGGTCGAAT TTATATCGTG AACGTAAAAA GAATCACCTT

                                121
Bm-osm-9_Bm1711.1                 TACAAATGGG TTGGCATGCG CAGTGGTGGT GAACTGTTGG CTGCTTTTGA AAAGGAAGGT GAAGATGGAG TTTTAAAGTT TGCGAACGAA AAGCTTATTT CAATGATGTA TGATGATGGT
Bm-osm-9_clone                    TACAAATGGG TTGGCATGCG CAGTGGTGGT GAACTGTTGG CTGCTTTTGA AAAGGAAGGT GAAGATGGAG TTTTAAAGTT TGCGAACGAA AAGCTTATTT CAATGATGTA TGATGATGGT
i3_LQ_NR42497|c19820/f1p1/3020    TACAAATGGG TTGGCATGCG CAGTGGTGGT GAACTGTTGG CTGCTTTTGA AAAGGAAGGT GAAGATGGAG TTTTAAAGTT TGCGAACGAA AAGCTTATTT CAATGATGTA TGATGATGGT
i2_LQ_NR42497|c51951/f1p6/3009    TACAAATGGG TTGGCATGCG CAGTGGTGGT GAACTGTTGG CTGCTTTTGA AAAGGAAGGT GAAGATGGAG TTTTAAAGTT TGCGAACGAA AAGCTTATTT CAATGATGTA TGATGATGGT

                                241
Bm-osm-9_Bm1711.1                 GCATCACCGC AAATGATTCG TTTTACAGAC TATGCAAAAT GGAAAAAGAC AACTAATGTA CAGCTGGGTA AAACAGAATC AAACTCTGTT GGCCAATTTG GTAGCAAATT CCGTGAACAT
Bm-osm-9_clone                    GCATCACCGC AAATGATTCG TTTTACAGAC TATGCAAAAT GGAAAAAGAC AACTAATGTA CAGCTGGGTA AAACAGAATC AAACTCTGTT GGCCAATTTG GTAGCAAATT CCGTGAACAT
i3_LQ_NR42497|c19820/f1p1/3020    GCATCACCGC AAATGATTCG TTTTACAGAC TATGCAAAAT GGAAAAAGAC AACTAATGTA CAGCTGGGTA AAACAGAATC AAACTCTGTT GGCCAATTTG GTAGCAAATT CCGTGAACAT
i2_LQ_NR42497|c51951/f1p6/3009    GCATCACCGC AAATGATTCG TTTTACAGAC TATGCAAAAT GGAAAAAGAC AACTAATGTA CAGCTGGGTA AAACAGAATC AAACTCTGTT GGCCAATTTG GTAGCAAATT CCGTGAACAT

                                361
Bm-osm-9_Bm1711.1                 CTTGGTCAGT GGAGGTTGAA CAAACGTGGT GTGGAAGGTG AGACGATAAT ACACCTATTG CTGAATCGTG AGGAACCAAT GTGTAGTGAA ATTGCTAGAA TTTTAATAAC ACGTTATCCA
Bm-osm-9_clone                    CTTGGTCAGT GGAGGTTGAA CAAACGTGGT GTGGAAGGTG AGACGATAAT ACACCTATTG CTGAATCGTG AGGAACCAAT GTGTAGTGAA ATTGCTAGAA TTTTAATAAC ACGTTATCCA
i3_LQ_NR42497|c19820/f1p1/3020    CTTGGTCAGT GGAGGTTGAA CAAACGTGGT GTGGAAGGTG AGACGATAAT ACACCTATTG CTGAATCGTG AGGAACCAAT GTGTAGTGAA ATTGCTAGAA TTTTAATAAC ACGTTATCCA
i2_LQ_NR42497|c51951/f1p6/3009    CTTGGTCAGT GGAGGTTGAA CAAACGTGGT GTGGAAGGTG AGACGATAAT ACACCTATTG CTGAATCGTG AGGAACCAAT GTGTAGTGAA ATTGCTAGAA TTTTAATAAC ACGTTATCCA

                                481
Bm-osm-9_Bm1711.1                 GGCCTTGCTA ATGATATTTA CCTCGGCGAC GAAATGTTCG GTCAAAGTGC ATTACATTTA GCAATTGTTC ATGATGATTA TGAAACTGTT CATTTATTAC TGCAAAACTC AGCAGAAGTG
Bm-osm-9_clone                    GGCCTTGCTA ATGATATTTA CCTCGGCGAC GAAATGTTCG GTCAAAGTGC ATTACATTTA GCAATTGTTC ATGATGATTA TGAAACTGTT CATTTATTAC TGCAAAACTC AGCAGAAGTG
i3_LQ_NR42497|c19820/f1p1/3020    GGCCTTGCTA ATGATATTTA CCTCGGCGAC GAAATGTTCG GTCAAAGTGC ATTACATTTA GCAATTGTTC ATGATGATTA TGAAACTGTT CATTTATTAC TGCAAAACTC AGCAGAAGTG
i2_LQ_NR42497|c51951/f1p6/3009    GGCCTTGCTA ATGATATTTA CCTCGGCGAC GAAATGTTCG GTCAAAGTGC ATTACATTTA GCAATTGTTC ATGATGATTA TGAAACTGTT CATTTATTAC TGCAAAACTC AGCAGAAGTG

                                601
Bm-osm-9_Bm1711.1                 AATGCCCGGG CTTGCGGGAC ATTTTTTCTA CCTGAGAATC AAAAGACATC TCGCAAATCA ACTGATTACC AAGGATATGC CTATTATGGC GAATATCCGT TAGCATTTGC TGCTTGTTTT
Bm-osm-9_clone                    AATGCCCGGG CTTGCGGGAC ATTTTTTCTA CCTGAGAATC AAAAGACATC TCGCAAATCA ACTGATTACC AAGGATATGC CTATTATGGC GAATATCCGT TAGCATTTGC TGCTTGTTTT
i3_LQ_NR42497|c19820/f1p1/3020    AATGCCCGGG CTTGCGGGAC ATTTTTTCTA CCTGAGAATC AAAAGACATC TCGCAAATCA ACTGATTACC AAGGATATGC CTATTATGGC GAATATCCGT TAGCATTTGC TGCTTGTTTT
i2_LQ_NR42497|c51951/f1p6/3009    AATGCCCGGG CTTGCGGGAC A-TTTTTCTA CCTGAGAATC AAAAGACATC TCGCAAATCA ACTGATTACC AAGGATATGC CTATTATGGC GAATATCCGT TAGCATTTGC TGCTTGTTTT

                                721
Bm-osm-9_Bm1711.1                 GGCAATAAAG ATATCTATGA TTTACTGATA CAATATGGCG CTGATCCAAA TCTACAAGAT ATGTTCGGCA ACACCATTCT ACATATGTGT GTCATCAACT ATAGCAATTC AATGTACAGT
Bm-osm-9_clone                    GGCAATAAAG ATATCTATGA TTTACTGATA CAATATGGCG CTGATCCAAA TCTACAAGAT ATGTTCGGCA ACACCATTCT ACATATGTGT GTCATCAACT ATAGCAATTC AATGTACAGT
i3_LQ_NR42497|c19820/f1p1/3020    GGCAATAAAG ATATCTATGA TTTACTGATA CAATATGGCG CTGATCCAAA TCTACAAGAT ATGTTCGGCA ACACCATTCT ACATATGTGT GTCATCAACT ATAGCAATTC AATGTACAGT
i2_LQ_NR42497|c51951/f1p6/3009    GGCAATAAAG ATATCTATGA TTTACTGATA CAATATGGCG CTGATCCAAA TCTACAAGAT ATGTTCGGCA ACACCATTCT ACATATGTGT GTCATCAACT ATAGCAATTC AATGTACAGT

                                841
Bm-osm-9_Bm1711.1                 TATGCAGTAA GGCATTGGGC GAAACCCGCT GATCCAAACA TTGTAAATGC AGCTGGTCTG ACACCATTGA CGCTTGCAAC AAAACTTGGA CGAAAGGATA TCTTTGAGGA AATGCTGGAG
Bm-osm-9_clone                    TATGCAGTAA GGCATTGGGC GAAACCCGCT GATCCAAACA TTGTAAATGC AGCTGGTCTG ACACCATTGA CGCTTGCAAC AAAACTTGGA CGAAAGGATA TCTTTGAGGA AATGCTGGAG
i3_LQ_NR42497|c19820/f1p1/3020    TATGCAGTAA GGCATTGGGC GAAACCCGCT GATCCAAACA TTGTAAATGC AGCTGGTCTG ACACCATTGA CGCTTGCAAC AAAACTTGGA CGAAAGGATA TCTTTGAGGA AATGCTGGAG
i2_LQ_NR42497|c51951/f1p6/3009    TATGCAGTAA GGCATTGGGC GAAACCCGCT GATCCAAACA TTGTAAATGC AGCTGGTCTG ACACCATTGA CGCTTGCAAC AAAACTTGGA CGAAAGGATA TCTTTGAGGA AATGCTGGAG

                                961
Bm-osm-9_Bm1711.1                 TTGATGAAAG TCGAGTTTTG GAGATTCAGT GATATGACGT GTAGTGCATA TCCTCTTACA GCTTTGGATA CTATTCGCCC CGATGGCTCA ACAAATTATG ATTCAGCGTT GATGACTGTG
Bm-osm-9_clone                    TTGATGAAAG TCGAGTTTTG GAGATTCAGT GATATGACGT GTAGTGCATA TCCTCTTACA GCTTTGGATA CTATTCGCCC CGATGGCTCA ACAAATTATG ATTCAGCGTT GATGACTGTG
i3_LQ_NR42497|c19820/f1p1/3020    TTGATGAAAG TCGAGTTTTG GAGATTCAGT GATATGACGT GTAGTGCATA TCCTCTTACA GCTTTGGATA CTATTCGCCC CGATGGCTCA ACAAATTATG ATTCAGCGTT GATGACTGTG
i2_LQ_NR42497|c51951/f1p6/3009    TTGATGAAAG TCGAGTTTTG GAGATTCAGT GATATGACGT GTAGTGCATA TCCTCTTACA GCTTTGGATA CTATTCGCCC CGATGGCTCA ACAAATTATG ATTCAGCGTT GATGACTGTG

                               1081
Bm-osm-9_Bm1711.1                 ATAAATGGTA GTACCAGTGA ACATTTAGAT ATGATTGGTA GCGAAGTGAT CCAACGACTT CTTGCTGACA AATGGAAAGC TTTTGCATCA AGAAAATTAT TTGAAAGGCT TGGATTGCTC
Bm-osm-9_clone                    ATAAATGGTA GTACCAGTGA ACATTTAGAT ATGATTGGTA GCGAAGTGAT CCAACGACTT CTTGCTGACA AATGGAAAGC TTTTGCATCA AGAAAATTAT TTGAAAGGCT TGGATTGCTC
i3_LQ_NR42497|c19820/f1p1/3020    ATAAATGGTA GTACCAGTGA ACATTTAGAT ATGATTGGTA GCGAAGTGAT CCAACGACTT CTTGCTGACA AATGGAAAGC TTTTGCATCA AGAAAATTAT TTGAAAGGCT TGGATTGCTC
i2_LQ_NR42497|c51951/f1p6/3009    ATAAATGGTA GTACCAGTGA ACATTTAGAT ATGATTGGTA GCGAAGTGAT CCAACGACTT CTTGCTGACA AATGGAAAGC TTTTGCATCA AGAAAATTAT TTGAAAGGCT TGGATTGCTC

                               1201
Bm-osm-9_Bm1711.1                 ATTTTACATC TCATATTTTT ATGTTTTGTT GTTTATATGC GGCCGTCTGA ACCGGAGCGA CTTACGTATA AATTAATTGC AACAGAATGG AATGATTGGG TGCGGTTATG CTTTGAAATT
Bm-osm-9_clone                    ATTTTACATC TCATATTTTT ATGTTTTGTT GTTTATATGC GGCCGTCTGA ACCGGAGCGA CTTACGTATA AATTAATTGC AACAGAATGG AATGATTGGG TGCGGTTATG CTTTGAAATT
i3_LQ_NR42497|c19820/f1p1/3020    ATTTTACATC TCATATTTTT ATGTTTTGTT GTTTATATGC GGCCGTCTGA ACCGGAGCGA CTTACGTATA AATTAATTGC AACAGAATGG AATGATTGGG TGCGGTTATG CTTTGAAATT
i2_LQ_NR42497|c51951/f1p6/3009    ATTTTACATC TCATATTTTT ATGTTTTGTT GTTTATATGC GGCCGTCTGA ACCGGAGCGA CTTATGTATA AATTAATTGC AACAGAATGG AATGATTGGG TGCGGTTATG CTTTGAAATT
```

```
                                1321
Bm-osm-9_Bm1711.1               CTAACGATAG CCAGTTGTGT GTTCTTCGTT TTCTTTCAGC AGTTTAGCGA ACTAAGAACG CAAGGATTTT ATGGATATAT CAGAAATCTG AAAACGGCTC CAGCTAAAAT TGTATTTCTT
Bm-osm-9_clone                  CTAACGATAG CCAGTTGTGT GTTCTTCGTT TTCTTTCAGC AGTTTAGCGA ACTAAGAACG CAAGGATTTT ATGGATATAT CAGAAATCTG AAAACGGCTC CAGCTAAAAT TGTATTTCTT
i3_LQ_NR42497|c19820/f1p1/3020  CTAACGATAG CCAGTTGTGT GTTCTTCGTT TTCTTTCAGC AGTTTAGCGA ACTAAGAACG CAAGGATTTT ATGGATATAT CAGAAATCTG AAAACGGCTC CAGCTAAAAT TGTATTTCTT
i2_LQ_NR42497|c51951/f1p6/3009  CTAACGATAG CCAGTTGTGT GTTCTTCGTT TTCTTTCAAC AGTTTAGCGA ACTAAGAACG CAGGGATTTT ATGGATATAT CAGAAATCTG AAAACGGCTC CAGCTAAAAT TGTATTTCTT

                                1441
Bm-osm-9_Bm1711.1               GGTGCCAATA TATGTATTCT TATATGTGTA CCTTTTCGTA TATCGGGCAA TGTTCAAGTT GAAGAAGCTT TATTAGTCTT TTCATTACCG GGTTCTTGGA TATTTTTCCT TTTCTTTGCC
Bm-osm-9_clone                  GGTGCCAATA TATGTATTCT TATATGTGTA CCTTTTCGTA TATCGGGCAA TGTTCAAGTT GAAGAAGCTT TATTAGTCTT TTCATTACCG GGTTCTTGGA TATTTTTCCT TTTCTTTGCC
i3_LQ_NR42497|c19820/f1p1/3020  GGTGCCAATA TATGTATTCT TATATGTGTA CCTTTTCGTA TATCGGGCAA TGTTCAAGTT GAAGAAGCTT TATTAGTCTT TTCATTACCG GGTTCTTGGA TATTTTTCCT TTTCTTTGCC
i2_LQ_NR42497|c51951/f1p6/3009  GGTGCCAATA TATGTATTCT TATATGTGTA CCCTTTCGTA TATCGGGCAA TGTTCAAGTT GAAGAAGCTT TATTAGTCTT TTCATTACCG GGTTCTTGGA TATTTTTCCT TTTCTTTGCC

                                1561
Bm-osm-9_Bm1711.1               AGAAGTGCCA AATTAACGGG ACCATTTGTG CAAATGATTT ATAGTATGAT TGCTGGCGAT ATGATTCGCT TTGCTATCAT TTCCGCTATA TTTCTTGTCT CATTTTCGCA AGTATTTTAT
Bm-osm-9_clone                  AGAAGTGCCA AATTAACGGG ACCATTTGTG CAAATGATTT ATAGTATGAT TGCTGGCGAT ATGATTCGCT TTGCTATCAT TTCCGCTATA TTTCTTGTCT CATTTTCGCA AGTATTTTAT
i3_LQ_NR42497|c19820/f1p1/3020  AGAAGTGCCA AATTAACGGG ACCATTTGTG CAAATGATTT ATAGTATGAT TGCTGGCGAT ATGATTCGCT TTGCTATCAT TTCCGCTATA TTTCTTGTCT CATTTTCGCA AGTATTTTAT
i2_LQ_NR42497|c51951/f1p6/3009  AGAAGTGCCA AATTAACGGG ACCATTTGTG CAAATGATTT ATAGTATGAT TGCTGGCGAT ATGATTCGCT TTGCTATCAT TTCCGCTATA TTTCTTGTCT CATTTTCGCA AGTATTTTAT

                                1681
Bm-osm-9_Bm1711.1               TTTCTGGGAA AGGATATGCA CGTGAAGCAA GAGCTCAATC CGCTGAATCC AGACTACTGC GAAGTGAAAG GCTATGATAT CTTCACGTAT TCCTCATTTC TAGAAACATT TATTACACTG
Bm-osm-9_clone                  TTTCTGGGAA AGGATATGCA CGTGAAGCAA GAGCTCAATC CGCTGAATCC AGACTACTGC GAAGTGAAAG GCTATGATAT CTTCACGTAT TCCTCATTTC TAGAAACATT TATTACACTG
i3_LQ_NR42497|c19820/f1p1/3020  TTTCTGGGAA AGGATATGCA CGTGAAGCAA GAGCTCAATC CGCTGAATCC AGACTACTGC GAAGTGAAAG GCTATGATAT CTTCACGTAT TCCTCATTTC TAGAAACATT TATTACACTG
i2_LQ_NR42497|c51951/f1p6/3009  TTTCTGGGAA AGGATATGCA CGTGAAGCAA GAGCTCAATC CGCTGAATCC AGACTACTGC GAAGTGAAAG GCTATGATAT CTTCACGTAT TCCTCATTTC TAGAAACATT TATTACACTG

                                1801
Bm-osm-9_Bm1711.1               TTTCGTGCTA GCATGGGTGG CTATGATTAT GAGGAATTTT CTTGCGCTAA TTACGAAGTG TTAACAAAAA TATTATTCGT ACTTTATATG TTTATAATGC CGATCATGTT GATTAATATA
Bm-osm-9_clone                  TTTCGTGCTA GCATGGGTGG CTATGATTAT GAGGAATTTT CTTGCGCTAA TTACGAAGTG TTAACAAAAA TATTATTCGT ACTTTATATG TTTATAATGC CGATCATGTT GATTAATATA
i3_LQ_NR42497|c19820/f1p1/3020  TTTCGTGCTA GCATGGGTGG CTATGATTAT GAGGAATTTT CTTGCGCTAA TTACGAAGTG TTAACAAAAA TATTATTCGT ACTTTATATG TTTATAATGC CGATCATGTT GATTAATATA
i2_LQ_NR42497|c51951/f1p6/3009  TTTCGTGCTA GCATGGGTGG CTACGATTAT GAGGAATTTT CTTGCGCTAA TTACGAAGTG TTAACAAAAA TATTATTCGT ACTTTATATG TTTATAATGC CGATCATGTT GATTAATATA

                                1921
Bm-osm-9_Bm1711.1               CTGATCGCCA TGATGGGTAA TACTTATACA ACAGTCATCG CACAAGCTGA AAAAGCTTGG AGGCAACAGT ACGCGCAAAT TGTAATGGTA TTGGAAAGAT CGGTG-AAAA AGGAAAAATT
Bm-osm-9_clone                  CTGATCGCCA TGATGGGTAA TACTTATACA ACAGTCATCG CACAAGCTGA AAAAGCTTGG AGGCAACAGT ACGCGCAAAT TGTAATGGTA TTGGAAAGAT CGGTG-AAAA AGGAAAAATT
i3_LQ_NR42497|c19820/f1p1/3020  CTGATCGCCA TGATGGGTAA TACTTATACA ACAGTCATCG CACAAGCTGA AAAAGCTTGG AGGCAACAGT ACGCGCAAAT TGTAATGGTA TTGGAAAGAT CGGTGAAAAA AGGAAAAATT
i2_LQ_NR42497|c51951/f1p6/3009  CTGATCGCCA TGATGGGTAA TACTTATACA ACAGTCATCG CACAAGCTGA AAAAGCTTGG AGGCAACAGT ACGCGCAAAT TGTAATGGTA TTGGAAAGAT CGGTG-AAAA AGGAAAAATT

                                2041
Bm-osm-9_Bm1711.1               AGCAGCGTGT CAATTAGAGT ATAGTATTAG ACTTAATGAA GCTAATGATG CTGGTATGGA AATACGTGGT CTAATGGTCA TCAAACAGAC AAAGAAAACT CGAGCAAGAC AAAGGAAGCA
Bm-osm-9_clone                  AGCAGCGTGT CAATTAGAGT ATAGTATTAG ACTTAATGAA GCTAATGATG CTGGTATGGA AATACGTGGT CTAATGGTCA TCAAACAGAC AAAGAAAACT CGAGCAAGAC AAAGGAAGCA
i3_LQ_NR42497|c19820/f1p1/3020  AGCAGCGTGT CAATTAGAGT ATAGTATTAG ACTTAATGAA GCTAATGATG CTGGTATGGA AATACGTGGT CTAATGGTCA TCAAACAGAC AAAGAAAACT CGAGCAAGAC AAAGGAAGCA
i2_LQ_NR42497|c51951/f1p6/3009  AGCAGCATGT CAATTAGAGT ATAGTATTAG ACTTAATGAA GCTAATGATG CTGGTATGGA AATACGTGGT CTAATGGTCA TCAAACAGAC AAAGAAAACT CGAGCAAGAC AAAGGAAGCA

                                2161
Bm-osm-9_Bm1711.1               AGCTATCACT AATTGGAAGA CAATTGGTCG AAAAGTAATT CATACTGTGG AAAGGTTGGG TGTTGATTAT GCACAGGAAT TACTTCATTC ATACAACTGT TTAATTGATG AACCAGCTGG
Bm-osm-9_clone                  AGCTATCACT AATTGGAAGA CAATTGGTCG AAAAGTAATT CATACTGTGG AAAGGTTGGG TGTTGATTAT GCACAGGAAT TACTTCATTC ATACAACTGT TTAATTGATG AACCAGCTGG
i3_LQ_NR42497|c19820/f1p1/3020  AGCTATCACT AATTGGAAGA CAATTGGTCG AAAAGTAATT CATACTGTGG AAAGGTTGGG TGTTGATTAT GCACAGGAAT TACTTCATTC ATACAACTGT TTAATTGATG AACCAGCTGG
i2_LQ_NR42497|c51951/f1p6/3009  AGCTATCACT AACTGGAAGA CAATTGGTCG AAAAGTAATT CATACTGTGG AAAGGTTGGG TGTTGGTTAT GCACAGGAAT TACTTCATTC ATACAACTGT TTAATTGATG AACCAGCTGG

                                2281
Bm-osm-9_Bm1711.1               TGCTGTAATA CTTAGACGAG ATACTGTATT TCCACCACCA ACAAGGTCTA TGACAAGATC ACAAGCTCAT GAAACCTCAA CAAATACGCA AAATGAAGAA AAAATCGAAC ATTTAAATAA
Bm-osm-9_clone                  TGCTGTAATA CTTAGACGAG ATACTGTATT TCCACCACCA ACAAGGTCTA TGACAAGATC ACAAGCTCAT GAAACCTCAA CAAATACGCA AAATGAAGAA AAAATCGAAC ATTTAAATAA
i3_LQ_NR42497|c19820/f1p1/3020  TGCTGTAATA CTTAGACGAG ATACTGTATT TCCACCACCA ACAAGGTCTA TGACAAGATC ACAAGCTCAT GAAACCTCAA CAAATACGCA AAATGAAGAA AAAATCGAAC ATTTAAATAA
i2_LQ_NR42497|c51951/f1p6/3009  TGCTGTAATA CTTAGACGAG ATACTGTATT TCCACCACCA ACAAGGTCTA TGACAAGATC ACGAGCTCAT GAAACCTCAA CAAATACGCA AAATGAAGAA AAAATCGAAC ATTTAAATAA

                                2401
Bm-osm-9_Bm1711.1               TATCTCAATC GAGAAAGAAA TAATCGTAAA TAATTGTTCC AACCAAACGT CTCTCCAATG TCAGTCACCG CGAGATTTTA ---------- ---------- ---------- ----------
Bm-osm-9_clone                  TATCTCAATC GAGAAAGAAA TAATCGTAAA TAATTGTTCC AACCAAACGT CTCTCCAATG TCAGTCACCG CGAGATTTTA AATTAATAAC AACCAAACAA AGCACAACTA TGATTCTGGA
i3_LQ_NR42497|c19820/f1p1/3020  TATCTCAATC GAGAAAGAAA TAATCGTAAA TAATTGTTCC AACCAAACGT CTCTCCAATG TCAGTCACCG CGAGATTTTA AATTAATAAC AACCAAACAA AGCACAACTA TGATTCTGGA
i2_LQ_NR42497|c51951/f1p6/3009  TATCTCAATC GAGAAAGAAA TAATCGTAAA TAATTGCTCC AACCAAACGT CTCTCCAATG TCAGTCACCG CGAGATTTTA AATTAATAAC AACCGAACAA AGCACAACTA TGATTCTGGA

                                2521
Bm-osm-9_Bm1711.1               -AAGTTGCCA ATGCGGAAAA CGGCACGAGG ACATCAGTTG GTTCCATCGT TAGACATACC GAATATGCCA GCTGCTGGAA CACCACCACG AGCAGTATCA CCGAGGATTC GAAAAGATAT
Bm-osm-9_clone                  GAAGTTGCCA ATGCGGAAAA CGGCACGAGG ACATCAGTTG GTTCCATCGT TAGACATACC GAATATGCCA GCTGCTGGAA CACCACCACG AGCAGTATCA CCGAGGATTC GAAAAGATAT
i3_LQ_NR42497|c19820/f1p1/3020  GAAGTTGCCA ATGCGGAAAA CGGCACGAGG ACATCAGTTG GTTCCATCGT TAGACATACC GAATATGCCA GCTGCTGGAA CACCACCACG AGCAGTATCA CCGAGGATTC GAAAAGATAT
i2_LQ_NR42497|c51951/f1p6/3009  GAAGTTGCCA ATGCGGAAAA CGGCACGAGG ACATCAGTTG GTTCCATCGT TAGACATACC GAATATGCCA GCTGCTGGAA CACCACCACG AGCAATATCA CCGATGATTC GAAAAGATAT
```

```
                                        2641
Bm-osm-9_Bm1711.1                       GTTTCGGCGC AAAGACAGTT TC-TGTGCGA GTTCATCATC TGCCAGCTCC AGTACGCAGG TTTGACATCA AGGATAAATA ATACGTCAAA AAATGTAACA ATTGCAGTCA ACTTCAATTT
Bm-osm-9_clone                          GTTTCGGCGC AAAGACAGTT TC-TGTGCGA GTTCATCATC TGCCAGCTCC AGTACGCAGG TTTGACATCA AGGATAAATA ATACGTCAAA AAATGTAACA ATTGCAGTCA ACTTCAATTT
i3_LQ_NR42497|c19820/f1p1/3020          GTTTCGGCGC AAAGACAGTT TCTTGTGCGA GTTCATCATC TGCCAGCTCC AGTACGCAGG TTTGACATCA AGGATAAATA ATACGTCAAA AAATGTAACA ATTGCAGTCA ACTTCAATTT
i2_LQ_NR42497|c51951/f1p6/3009          GTTTCGGCGC AAAGACAGTT TC-TGTGCGA GTTCATCATC TGCCAGCTCC AGTACGCAGG TTTGACATCA AGGATAAATA ATACGTCAAA AAATGTAACA GTTGCAGTCA ACTTCAATTT

                                        2761
Bm-osm-9_Bm1711.1                       CAGTGGCTGA ---------- ---------- ---------- ---------- ---------- ---------- ---------- ---------- ----
Bm-osm-9_clone                          CAGTGGCTGA ---------- ---------- ---------- ---------- ---------- ---------- ---------- ---------- ----
i3_LQ_NR42497|c19820/f1p1/3020          CAGTGGCTGA ATTTATGATG TGTTCTCAAA ATTGTTTTAA TTAC-ATTCT TCTAATGCCA GTTTGTTTGC CTTATCAATA AAAATTTACG ----
i2_LQ_NR42497|c51951/f1p6/3009          CAGTGGCTGA ATTTATGATG TGTTCTCAAA ATTGTTTTAA TTACAATTCT TCTAATGCCA G-TTGTTTGC CTGATC-ATA AAAATTTACG AGAC
```
