## Supplementary material for "Genetic and functional diversification of chemosensory pathway receptors in mosquito-borne filarial nematodes": S8 Figure

```
                      1
inactive_FBtr0070969  MKFLLKKCLR KKAPE----- -MKPGAILDA VISQSSATAC KCLLYKLADY KRGGDLIDAI NSGGLIAVEQ LIREQFGVFM YNDGKG-QVI NRAEFLRWKY RDHTEVTIPI EASLSIHDPL GKWEDHKACW
Ce-osm-9_B0212.5.1    MGGGSSRNKT EPRGEGV-KL AFDP----DE KWSNLYRERE KNHLYKWVAV RKGGELINIY ERDGEEGVLK FAEEKLLTIL YDEGQNPKLV TYSDYIKWK- -KGVNVQLGL SE-ESVDMQQ SRFKEHYALW
Bm-osm-9_clone        MGQLKSKILH QTGGEDIDNL KEDP----DD QWSNLYRERK KNHLYKWVGM RSGGELLAAF EKEGEDGVLK FANEKLISMM YDDGASPQMI RFTDYAKWK- -KTTNVQLGK TESNSVGQFG SKFREHLGQW
Bm-osm-9_Bm1711.1     MGQLKSKILH QTGGEDIDNL KEDP----DD QWSNLYRERK KNHLYKWVGM RSGGELLAAF EKEGEDGVLK FANEKLISMM YDDGASPQMI RFTDYAKWK- -KTTNVQLGK TESNSVGQFG SKFREHLGQW

                      131
inactive_FBtr0070969  QMQYRGALGE SLLHVLIICD SKVHTKLARV LLRVFPNLAL DVMEGEEYLG ASALHLSIAY SNNELVADLI EAGADIHQRA IGSFFLPRDQ QRANPAKSTD YEGLAYMGEY PLAWAACCAN ESVYNLLVDC
Ce-osm-9_B0212.5.1    KLNKRGVEGE NLIHLLLNRE QQVCYEIARI LLKRFPGMAN DIYLGDEQFG QSALHLAIVH DDYETVSLLL NSKADVNARA CGNFFLPEDF KLTN--KITD YQGYAYYGEY PLAFAACFGN KDIYDLLIQF
Bm-osm-9_clone        RLNKRGVEGE TIIHLLLNRE EPMCSEIARI LITRYPGLAN DIYLGDEMFG QSALHLAIVH DDYETVHLLL QNSAEVNARA CGTFFLPENQ KTSR--KSTD YQGYAYYGEY PLAFAACFGN KDIYDLLIQY
Bm-osm-9_Bm1711.1     RLNKRGVEGE TIIHLLLNRE EPMCSEIARI LITRYPGLAN DIYLGDEMFG QSALHLAIVH DDYETVHLLL QNSAEVNARA CGTFFLPENQ KTSR--KSTD YQGYAYYGEY PLAFAACFGN KDIYDLLIQY

                      261
inactive_FBtr0070969  GSDPDAQDSF GNMILHMVVV CDKLDMFGYA LRHPKTPAKN GIVNQTGLTP LTLACKLGRA EVFREMLELS AREFWRYSNI TCSGYPLNAL DTLLPDGRTN WNSALFIILN GTKPEHLDML DGGIIQRLLE
Ce-osm-9_B0212.5.1    GANPNLQDSF GNTILHMCVI NYSSSMYSYA VRHWAKPADP HVVNHAGFTP LTLATKLGRK QIFEEMLEIM KVEFWRFSDM TCSAYPLNTL DTIQPDGSTN YDSALMTVIN GSTPEHLDMI GSEVIQRLLA
Bm-osm-9_clone        GADPNLQDMF GNTILHMCVI NYSNSMYSYA VRHWAKPADP NIVNAAGLTP LTLATKLGRK DIFEEMLELM KVEFWRFSDM TCSAYPLTAL DTIRPDGSTN YDSALMTVIN GSTSEHLDMI GSEVIQRLLA
Bm-osm-9_Bm1711.1     GADPNLQDMF GNTILHMCVI NYSNSMYSYA VRHWAKPADP NIVNAAGLTP LTLATKLGRK DIFEEMLELM KVEFWRFSDM TCSAYPLTAL DTIRPDGSTN YDSALMTVIN GSTSEHLDMI GSEVIQRLLA

                      391
inactive_FBtr0070969  EKWKTFAQNQ FLKRLLILST HLLCLSVSVY LRPAHDGEAE DE--DSEGSD ASAAALLDIQ SDEGDSGGGD YNAQTVARYC AEFATLVGVL SYVIFQQGDE IKNQGLSAFL KQLSHAPAKA IFLFSNLLIL
Ce-osm-9_B0212.5.1    DKWKAFAQRK LIERLVLLIV QLITLSIVVY IRPTELPRLY ME--DPQWDD ---------- ---------- -----YIRTA CELLTILNCI FFVGYQQLGE IRTQGMRGYL RNLKTAPAKA VFCIANLFLL
Bm-osm-9_clone        DKWKAFASRK LFERLGLLIL HLIFLCFVVY MRPSEPERLT YKLIATEWND ---------- ---------- -----WVRLC FEILTIASCV FFVFFQQFSE LRTQGFYGYI RNLKTAPAKI VFLGANICIL
Bm-osm-9_Bm1711.1     DKWKAFASRK LFERLGLLIL HLIFLCFVVY MRPSEPERLT YKLIATEWND ---------- ---------- -----WVRLC FEILTIASCV FFVFFQQFSE LRTQGFYGYI RNLKTAPAKI VFLGANICIL

                      521
inactive_FBtr0070969  ACIPFRLIGD TDTEEAILIF AVPGSWFLLM FFAGAIRLTG PFVTMIYSMI TGDMFTFGII YCIVLCGFSQ AFYFLYK--- ---------- GHPQVQSTMF NTYTS---TW MALFQTTLGD YNYPDLNQTT
Ce-osm-9_B0212.5.1    LCIPFRLMKK HEIEEALFVF ALPGSWIFLL FFARSAKLTG PFVQMIYSMI AGDMIRFAII SAIFLVSFSQ VFYFVGKDMD AKQKLEDTNP HACRISGYTI YTYNTFPETF ITLFRASMGG YDYEEFSCAN
Bm-osm-9_clone        ICVPFRISGN VQVEEALLVF SLPGSWIFFL FFARSAKLTG PFVQMIYSMI AGDMIRFAII SAIFLVSFSQ VFYFLGKDMH VKQELNPLNP DYCEVKGYDI FTYSSFLETF ITLFRASMGG YDYEEFSCAN
Bm-osm-9_Bm1711.1     ICVPFRISGN VQVEEALLVF SLPGSWIFFL FFARSAKLTG PFVQMIYSMI AGDMIRFAII SAIFLVSFSQ VFYFLGKDMH VKQELNPLNP DYCEVKGYDI FTYSSFLETF ITLFRASMGG YDYEEFSCAN

                      651
inactive_FBtr0070969  YPNLSKTVFV IFMIFVPILL LNMLIAMMGN TYVTVIEQSE KEWMKQWAKI VVTLERAVPQ ADAKGYLEAY SIPLGPSDDS GFEVRGVMVI KSKSKTRAKQ RKGAVSNWKR VGRVTLTALK KRGMTGEEMR
Ce-osm-9_B0212.5.1    YQALTKTLFV LYMFVMPIMM INILIAMMGN TYTTVIAQAE KAWRQQYAQI VMVLERSVGK ERLAASQLEY SIRLDQEGSS GMEVRGLMVI KQTKKTRARQ RKQAIYNWKT IGRKVIHTID KVGT--EQAV
Bm-osm-9_clone        YEVLTKILFV LYMFIMPIML INILIAMMGN TYTTVIAQAE KAWRQQYAQI VMVLERSVKK EKLAACQLEY SIRLNEANDA GMEIRGLMVI KQTKKTRARQ RKQAITNWKT IGRKVIHTVE RLGV--DYAQ
Bm-osm-9_Bm1711.1     YEVLTKILFV LYMFIMPIML INILIAMMGN TYTTVIAQAE KAWRQQYAQI VMVLERSVKK EKLAACQLEY SIRLNEANDA GMEIRGLMVI KQTKKTRARQ RKQAITNWKT IGRKVIHTVE RLGV--DYAQ

                      781
inactive_FBtr0070969  RLMWGRASIS S--------- -----PVKVT KQKLKDPYNL HTDSDFTNAM DMLTFASNPA SSNGVTLRS- -------VTA PPPAP----- ----PAPDPF RELIMMSDQR PETHDPHYFA GLQQLANKA-
Ce-osm-9_B0212.5.1    LLLHGHDRLD R-------VY EDHVQPEKVP SRS-RTPTRI GTTLNSSKRL KTTTMVVGAA VTNTHEVRTD EAVNSMLLSA PPSLSGEGAT MDWQPSITPV EERSESKSQE DRSEASTPNL GIHRTTPKAD
Bm-osm-9_clone        ELLHSYNCLI DEPAGAVILR RDTVFPP--P TRS-MTRSQA HETSTNTQNE EKIEHLNNIS IEKEIIVNNC SNQTSLQCQS PRDFKLITTK ---------- ---------- QSTTMI---- ----------
Bm-osm-9_Bm1711.1     ELLHSYNCLI DEPAGAVILR RDTVFPP--P TRS-MTRSQA HETSTNTQNE EKIEHLNNIS IEKEIIVNNC SNQTSLQCQS PRDFKVANAE NGTRTSVGSI VRHTEYASCW NTTTSS---- ----------

                      911
inactive_FBtr0070969  --FDLVEQTM KTQPQAPVAK KVDPLPVASV AKASPAAPAT QATATAAA-- -------ASD LMAMPLPISN LSNLFQDPKD IVDPKKLEEF MAMLAEVETE ESDSGGPILG KLSLAKRTHN ALSKAEIRRD
Ce-osm-9_B0212.5.1    SPIRVVEYSR TIRVRAADTI PSIELPNIPT KQTSSTPPHR AVSPRLRADM FRRHQQPASF DQSPPLPPTN ---------- ---------- ---------- ---------- ---------- --DKSE----
Bm-osm-9_clone        ----LEKLPM RKTARGHQLV PSLDIPNMP- --AAGTPP-R AVSPRIRKDM FRRKDSFCAS SSS-----AS ---------- ---------- ---------- ---------- ---------- --SSTQV---
Bm-osm-9_Bm1711.1     ----ITEDSK RYVSAQRQFL CEFIICQLQ- --YAGLTS-R ---------- --------IN NTS-----KN ---------- ---------- ---------- ---------- ---------- --VTIAVN--

                      1041
inactive_FBtr0070969  QQGFEGHSHG QFQPMSSVWA PPGLDVDTGF HFDEAVAEEV LTIEQEAEVE TEDGNGGQDS EDIPTAEEVH ATMKQFHLRK CQPAQDEAAR RAKSARVRRR NKVSPEQSDD PDERSQRGRS AYTRRTQSPP
Ce-osm-9_B0212.5.1    ---------- ---------- ---------- ---------- ---------- ---------- ---------- ---------- ---------- ---------- ---------- ---------- ----------
Bm-osm-9_clone        ---------- ---------- ---------- ---------- ---------- ---------- ---------- ---------- ---------- ---------- ---------- ---------- ----------
Bm-osm-9_Bm1711.1     ---------- ---------- ---------F NFSG------ ---------- ---------- ---------- ---------- ---------- ---------- ---------- ---------- ----------

                      1171
inactive_FBtr0070969  DPLEPWSTRE LQDINKILAR K
Ce-osm-9_B0212.5.1    ---------- ---------- -
Bm-osm-9_clone        ---------- ---------- -
Bm-osm-9_Bm1711.1     ---------- ---------- -
```
