## Supplementary material for "Genetic and functional diversification of chemosensory pathway receptors in mosquito-borne filarial nematodes": S9 Figure

```
                     1
Ce-tax-4_ZC84.2.1    MSTAEPAPDP TNPSTSGLAP TTNGIGSPPP TASAATKFSI LTKFLRR--- ---------- --KNQVHTTT AQQNEFMQKY MPNGNSNAVQ PAATGGQPAS SDGGSAIEVP PPKESYAVRI RKYLANYTQD
Bm-tax-4_Bm7343.1    MFSK--SHDD T--YENHLKK REN------LP NASAMKNISN NDSFQKRHKS LRSNKICNIN EEENQAPSTV TNTPQLNSID SNKSCEKAKQ SLIETSAPTS LTSKQPATEA PSATTHETKW QYLLNKWVLD
Bm-tax-4_clone       MFSK--SHDD T--YENHLKK REN------LP NASAMKNISN NDSFQKRHKS LRSNKICNIN EEENQAPSTV TNTPQLNSID SNKSCEKAKQ SLIETSAPTS LTSKQPATEA PSATTHETKW QYLLNKWVLD

                   131
Ce-tax-4_ZC84.2.1    PSTDNFYYWT CVVTVAYIYN LLFVIARQVF NDLIGPSSQS LCRFYNGTLN STTQVECTYN MLTNMKEMPT YSQYPDLGWS KYWHFRMLWV FFDLLMDCVY LIDTFLNYRM GYMDQGLVVR EAEKVTKAYW
Bm-tax-4_Bm7343.1    TKDEFYYYWL SIVSCAFTYN LIVVIGKLFF ---------- ---------- ---------- -------LLS RSVFNDLAYG YYW---IAWL LVDFIMDITY VLDMFVRSRT GFLEQGLLVR DISRISKLYL
Bm-tax-4_clone       TKDEFYYYWL SIVSCAFTYN LIVVI----- ---------- ---------- ---------- ---------A RSVFNDLAYG YYW---IAWL LVDFIMDITY VLDMFVRSRT GFLEQGLLVR DISRISKLYL

                   261
Ce-tax-4_ZC84.2.1    QSKQYRIDGI SLIPLDYILG WPIPYINWRG LPILRLNRLI RYKRVRNCLE RTETRSSMPN AFRVVVVVWY IVIIIHWNAC LYFWISEWIG LGTDAWVYGH LNKQSLPDDI TDTLLRRYVY SFYWSTLILT
Bm-tax-4_Bm7343.1    KSLQFKLDII SVLPFDLILS ----FIFQRS IPYLRFNRII RYPRFSDFVD RTETRSSMPN AFRIFCVIVN IIVIIHWNAC IYFFISEMIG LGSDGWVYGP LNKQSLPDGV EDTLVRRYIY SFYWSTLILT
Bm-tax-4_clone       KSLQFKLDII SVLPFDLILS ----LIFQRS IPYLRFNRII RYPRFSDFVD RTETRSSMPN AFRIFCVIVN IIVIIHWNAC IYFFISEMIG LGSDGWVYGP LNKQSLPDGV EDTLVRRYIY SFYWSTLILT

                   391
Ce-tax-4_ZC84.2.1    TIGEVPSPVR NIEYAFVTLD LMCGVLIFAT IVGNVGSMIS NMSAARTEFQ NKMDGIKQYM ELRKVSKQLE IRVIKWFDYL WTNKQSLSDQ QVLKVLPDKL QAEIAMQVHF ETLRKVRIFQ DCEAGLLAEL
Bm-tax-4_Bm7343.1    TIGEVPSPKR NIEFLFVIMD LMCGVLIFAT IVGNVGSAIS NMSLARTKFQ NKMDGIKQYM KLRKVNKELE TRVMKWFDYL WEHKQSLSDQ RVLKVLPDKL QTEIAMQVHY ETLRRVRIFQ DCEAGLLAEL
Bm-tax-4_clone       TIGEVPSPKR NIEFLFVIMD LMCGVLIFAT IVGNVGSAIS NMSLARTKFQ NKMDGIKQYM KLRKVNKELE TRVMKWFDYL WEHKQSLSDQ RVLKVLPDKL QTEIAMQVHY ETLRRVRIFQ DCEAGLLAEL

                   521
Ce-tax-4_ZC84.2.1    VLKLQLQVFS PGDFICKKGD IGREMYIVKR GRLQVVDDDG KKVFVTLQEG SVFGELSILN IAGSKNGNRR TANVRSVGYT DLFVLSKTDL WNALREYPDA RKLLLAKGRE ILKKDNLLDE NAPEEQKTVE
Bm-tax-4_Bm7343.1    VLKLQQQIFS PGDYICKKGD IGREMYIVKR GKLQVVADDG IKVFATLQEG AVFGELSILN IAGSKNGNRR TANVRSVGYT DLFALNKNDL WTALKEYPDA RKLLIAKGRE ILRKDGLLDE DAPEEQMTAE
Bm-tax-4_clone       VLKLQQQIFS PGDYICKKGD IGREMYIVKR GKLQVVADDG IKVFATLQEG AVFGELSILN IAGSKNGNRR TANVRSVGYT DLFALNKNDL WTALKEYPDA RKLLIAKGRE ILRKDGLLDE DAPEEQMTAE

                   651
Ce-tax-4_ZC84.2.1    EIAEHLNNAV KVLQTRMARL IVEHSSTEGK LMKRIEMLEK HLSRYKALAR RQKTMHGVSI DGGDISTDGV DERVRPPRLR QTKTIDLPTG TESESLLK
Bm-tax-4_Bm7343.1    EMAKNLQNTL KIMQTKMARF AAEFSSVKTK LLARIEYLET QLAKYQI--- ---------- --NDNSTSSN D--------- -----DYQMI CDS-----
Bm-tax-4_clone       EMAKNLQNTL KIMQTKMARF AAEFSSVKTK LLARIEYLET QLAKYQI--- ---------- --NDNSTSSN D--------- -----DYQMI CDS-----
```
