## Supplementary material for "Genetic and functional diversification of chemosensory pathway receptors in mosquito-borne filarial nematodes": S10 Figure

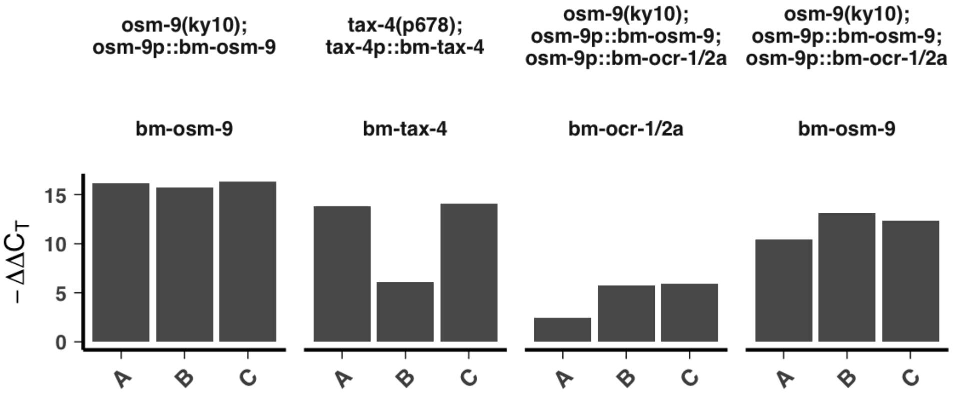


| | Target | | | |
|---|---|---|---|---|
| **Genotype** | **Y45F10D.4** | **bm-osm-9** | **bm-tax-4** | **bm-ocr-1/2a** |
| **N2** | 24.18 | ND | ND | ND |
| **osm-9(ky10)** | 23.93 | ND | | ND |
| **osm-9(ky10); osm-9p::bm-osm-9** | | | | |
| | 24 | 23.88 | | |
| | 24.14 | 24.44 | | |
| | 23.72 | 23.47 | | |
| **osm-9(ky10); osm-9p::bm-osm-9; osm-9p::bm-ocr-1/2a** | | | | |
| | 23.56 | 29.19 | | 37.20 |
| | 23.96 | 26.91 | | 34.30 |
| | 23.79 | 27.51 | | 33.90 |
| **tax-4(p678)** | 23.54 | | ND | |
| **tax-4(p678); tax-4p::bm-tax-4** | | | | |
| | 24.3 | | 26.98 | |
| | 23.99 | | 34.38 | |
| | 23.72 | | 26.13 | |

*Note:*
ND = Not Detected
